# Flexible pivoting of dynamin PH-domain catalyzes fission: Insights into molecular degrees of freedom

**DOI:** 10.1101/531814

**Authors:** K. K. Baratam, K. Jha, A. Srivastava

## Abstract

The neuronal dynamin1 functions in the release of synaptic vesicles by orchestrating the process of GTPase-dependent membrane fission. Dynamin1 associates with the plasma membrane-localized phosphatidylinositol-4,5-bisphosphate (PIP2) with its centrally-located pleckstrin homology domain (PHD). The PHD is dispensable as fission can be managed, albeit at much slower rates, even when the PHD-PIP2 interaction is replaced by a generic polyhistidine- or polylysine-lipid interaction. However, even when the PHD is present, the length of the dynamin scaffold and in turn the membrane remodeling and fission rates are severely restricted with mutations such as I533A on membrane-interacting variable loop 1 (VL1) of PHD. These observations suggest that PIP2-containing membrane interactions of PHD could have evolved to expedite fission to fulfill the requirement of rapid kinetics of synaptic vesicle recycling. Here, we use a suite of multiscale modeling approaches that combine atomistic molecular dynamics simulations, mixed resolution membrane mimetic models, coarse-grained molecular simulations and advanced free-energy sampling methods (metadynamics and umbrella sampling) to explore PHD-membrane interactions. Our results reveal that: (a) the binding of PHD to PIP2-containing membranes modulates the lipids towards fission-favoring conformations and softens the membrane, (b) that PHD engages another loop (VL4) for membrane association, which acts as an auxiliary pivot and modulates the orientation flexibility of PHD on the membrane – a mechanism we believe may be important for high fidelity dynamin collar assembly on the membrane. (c) Through analyses of our trajectories data and free-energy calculations on membrane-bound WT and mutant systems, we also identify key residues on multiple VLs that stabilizes PHD membrane association. And we suggest experiments to explore the ability of PHD to associate with membrane in orientations that favors faster fission. Together, these insights provide a molecular-level understanding of the “catalytic” role of the PHD in dynamin-mediated membrane fission.

**SIGNIFICANCE:** Dynamin, a large multi-domain GTPase, remodels the membrane by self-assembling onto the neck of a budding vesicle and induces fission by its energy driven conformational changes. In this work, we use multi-scale molecular simulations to probe the role of dynamin’s pleckstrin-homology domain (PHD), which facilitates membrane interactions. Notably, PHD is dispensable for fission as is the case with extant bacterial and mitochondrial dynamins. However, reconstitution experiments suggest that the functional role of PHD in neuronal-membrane goes beyond that of an adaptor domain as it possibly ‘expedites’ the fission reaction during synaptic vesicle recycling. We provide a molecular-dynamics picture of how PHDs make membranes more pliable for fission and suggest new insights into the molecular-level processes driving the expedited fission behavior.

## INTRODUCTION

Dynamin is a multi-domain GTPase that self-assembles into a helical collar and catalyzes membrane fission leading to the release of nascent clathrin-coated vesicles during endocytosis (1–5). The GTPase domain (G-domain), bundle-signaling element (BSE) and the stalk domain are the three conserved domains in dynamin which are required to evoke stimulated GTPase activity upon self-assembly on the membrane (6). In addition to these domains, classical dynamins contain a proline rich domain (PRD) that interacts with SH3 domain-containing partner proteins. It also contains a pleckstrin-homology domain (PHD) using which it binds to plasma membrane-localized lipid phosphatidyinositol-4,5-bisphosphate (PIP2). GTP hydrolysis is proposed to orchestrate a series of conformational changes in the self-assembly to surmount the activation barrier to membrane fission (7, 8).

Structurally, the ubiquitous dynamin1 PHD has a C-terminal *α*-helix and a core *β*-sandwich with variable loops between the *β*-strands that form the binding pocket for PIP2. Crystal structure of the dynamin1 PHD (PDB: 1DYN) suggests a *β*-sandwich structure with two *β*-sheets, one with 4 and the other with 3 *β*-strands, oriented in an antiparallel arrangement. The structure defines 4 loops referred to as variable loops (VLs) as shown in Fig.1. These loops significantly differ from one another in hydrophobicity and electrostatics. VL1 (^531^IGIMKGG^537^) contains a hydrophobic stretch that inserts in the membrane and assists dynamin’s subcellular localization (9). VL3 (^590^NTEQRNVYKDY^600^) is highly polar and has also been shown to be important for membrane association (10, 11). While PHD helps classical dynamins to engage with the membrane(12, 13), numerous lines of evidence suggest that it functions more than a generic membrane anchor (3, 14–20). Genetic neurological disorders such as centronuclear myopathies and the Charcot-Marie-Tooth disease are linked to mutations in the PHD that map to regions distinct from those involved in membrane binding (21, 22). Cellular assays combined with biochemical and microscopic analysis of membrane fission with point mutants in the PHD of dynamin1 suggest that this domain might induce local membrane curvature by shallow insertion of one of its loops (17), and that such point mutations alter dynamics and orientation of the PHD on the membrane (18). Experimentally guided modeling has shown that tilting of the PHD could be responsible for creating a low energy pathway towards reaching the hemi-fission state (23). GTP hydrolysis is required to initiate fission through constriction (24) but whether this is adequate to achieve complete and leakage-free vesicle scission is still not clear (23, 25). Recent models indicates that the GTPase-driven scaffold constriction brings the membrane in close proximity but not enough to cross the energy barrier for fission and that stochastic fluctuations are responsible for the crossover from the constricted stage to the hemi-fission state (20, 26). Indeed, analysis of membrane fission at the single event resolution indicates that replacement of the PHD-PIP2 interaction with a generic lipid-protein interaction forms long-lived, highly constricted pre-fission intermediates on the membrane which manifests in a dramatic dampening of fission kinetics (3). Recent cryo-EM data of the dynamin polymer assembled on tubular membranes reveal complex and variable geometries of the PHD (23, 25, 27, 28). The variable PHD configurations may reflect an evolutionary requirement for generating membrane torque necessary for traversing a pathway whereby membrane constriction guarantees leakage-free fission as shown in some of the recent models (29).

**Figure 1:**
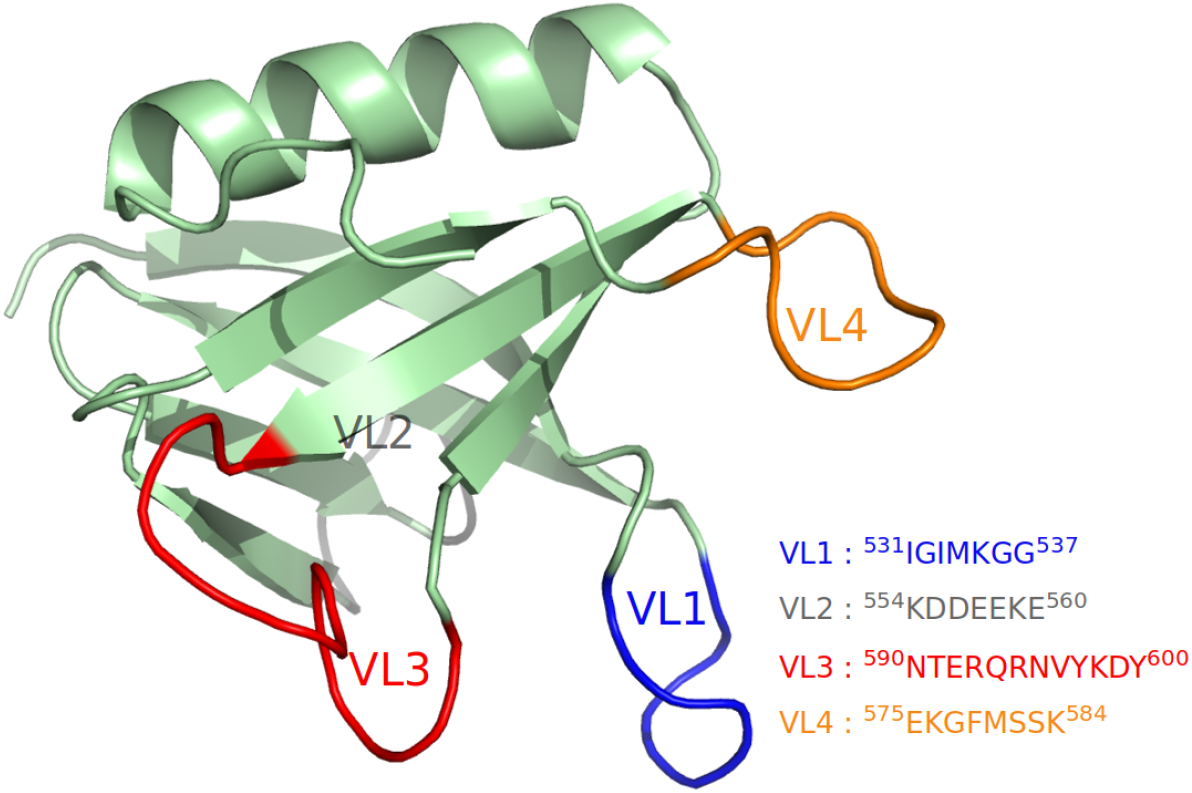
Pleckstrin Homology Domain(PHD) and its variable loops

Guided by these observations, and to circumvent the difficulty of observing short-lived transitions states of PHD-PIP2 interaction in an experimental scenario, we apply multi-scale molecular simulations to study the membrane association behavior of dyn-PHD in great molecular details. Undulation spectra extracted from Martini coarse-grained (CG) trajectories of large membrane systems (with and without PHD) were used to observe the effect on membrane physical properties. These methods also test the hypothesis of local curvature generation by the PHD. We then explore the conformational changes induced by PHD in lipids of the membrane using All-Atom molecular dynamics (AAMD) simulations. Mixed-resolution Highly Mobile Membrane-Mimetic Model (HMMM)-based molecular dynamics simulations initiated from different orientations were used to explore possible membrane association pathways. Membrane association geometries were deciphered using data from both AAMD simulations as well as by free energy difference calculations from enhanced sampling methods such as Metadynamics and Umbrella Sampling simulations. PHD is a ubiquitous membrane adaptor and PHD-membrane interactions have been studied thoroughly for several proteins in great details both in experiments and in molecular simulations (14, 30–42). In our work, we also look at the membrane association mechanism and intricate binding geometries and interactions of dyn-PHD in light of other available studies and highlight the similarities and differences while connecting the PHD-membrane interactions to dynamin-assisted fission process (3, 15, 17, 18, 23, 43, 44). Together, our results indicate that the presence of the PHD causes significant changes in lipid conformations and the membrane bending rigidity thus providing mechanistic insights into its catalytic effect on membrane fission due to dynamin. These effects are managed by flexible pivoting of the previously described variable loop1 (VL1) and the yet-unexplored variable loop (VL4) in the PHD with the membrane.

## MATERIALS AND METHODS

Complete description of the all-atom simulation parameters, metadynamics simulation parameters, umbrella simulations details as well as details of the Martini and HMMM CG modelling are described below or in the supporting information (SI). Moreover, all input files, including starting coordinates of the various systems, required to run all the simulation jobs for all methods, are made available at a Github location (https://github.com/codesrivastavalab/dynPHD).

### Coarse-grained Simulations

Two coarse-grained bilayer systems with and without dyn1-PHDs (randomly oriented) were generated using martini bilayer builder in CHARMM-GUI online server (45). The lipid bilayer was composed of 2048 lipids with a composition of DOPC-DOPS-PIP2 (80:19:1). These systems were simulated in NPT ensemble using MARTINI Polarizable force-field (Version 2.2P) in gromacs-5.1.2 (46–48). Both these systems were minimized using steepest descent algorithm followed by an equilibration of 5-10 ns before proceeding to 4 *μs* production run at 310 K temperature and 1 atm pressure. The temperature and pressure were maintained at 310 K and 1 atm using V-rescale thermostat and Berendsen barostat (with semi-isotropic coupling scheme) respectively. To understand the influence of dyn1-PHD on the mechanical properties of the lipid bilayer, we estimated the bending modulus (BM) of the lipid bilayer with and without dyn1-PHDs. In this regard we adopted the reciprocal space based method Brandt and co-workers(49) to obtain membrane undulation fluctuation spectra. The spectra so obtained was used to extract the BM using the Helfrich theory. To evaluate the mean and standard deviation on bending modulus estimate, we performed the spectra evaluation on the full trajectory as well as on blocks of 1 *μs* of the full trajectory. The reported values were evaluated from the estimates from individual blocks. All the relevant datafiles, python script and c++ codes can be found at the following github location (https://github.com/codesrivastavalab/dynPHD).

### HMMM Simulations

HMMM simulations (50, 51) were performed using gromacs-2019.1 with CHARMM36m protein force field (52) and the CHARMM36 phospholipid force field (53) with a time step of 2 fs. We ran 12 different replicates with starkly different initial orientations of PHD and each system was run for a minimum of 500 ns. The membrane was composed of 80:19:1 mixture of DOPC, DOPS and POPI headgroups (100 lipid molecules per leaflet). The core of the bilayer was made of DCLE molecules (~1200). The bilayer-PHD systems were solvated and ionizedwith 150 mM NaCl using VMD (54). For each system, the following simulation protocol was used: the system was equilibrated for 20 ns prior to a NVT simulation of 100 ns. Non-bonded forces were calculated with a 12 Å cutoff (10 Å switching distance). Long-range electrostatic forces were calculated at every other time step using the particle mesh Ewald method (55). The system is maintained at a temperature of 310 K using Langevin thermostat with friction coefficient of 0.5 *ps*^−1^.

### All-Atom Simulations

We performed series of AAMD NPT-simulations on WT and mutant dyn-PHD with membrane using gromacs-2019.1 (Table 1) to understand the dynamics and to obtain deeper molecular insights into dyn1-PHD on the membrane surface. Each of these systems were energy minimized and equilibrated for 20 ns. Each system comprises of PHD, ~ 200 lipids, ~ 23000 water molecules and 150 mM NaCl. CHARMM36 force-field (53) was used for particle definitions. Analysis of thickness variation across the membrane surface and protein and lipid conformational analysis were carried out using either an in-house VMD TCL script or VMD membrane plugin tool (56).

**Table 1:** List of of WT and mutant PHDs used for all-atom membrane simulations with membrane composition of DOPC-DOPS-PIP2 (80:19:1)

| Membrane+PHD variant | Simulation time (ns) |
| --- | --- |
| Wild Type | 500 |
| I533A | 500 |
| F579A | 500 |
| K539A | 500 |
| K583A | 500 |
| Y600L | 500 |
| I533A-F579A | 500 |
| F579K-K583A | 500 |
| F579A-S581K | 500 |
| K539A-K583A | 500 |

### Cryo-EM analysis

Cryo-EM map (28) for the constricted collar (K44A mutant) of dynamin (EMD-2701) was obtained from EM databank (www.emdatabank.org). The map is visualized using chimera (57) and is colored according to the cylinder radius. An in-house R code (58) was used to calculate the distribution of angles taken by PHDs in the Cryo-EM map. Our code is available on the aforementioned Github repository.

### Protein-Ligand Metadynamics

The X-ray solved structure of dyn1-PHD (protein) was obtained from protein data bank (1DYN) (59). The force-field parameters of inositol triphosphate (ligand) corresponding to CHARMM22 force-field were generated using Swissparam server (60). The initial configurations of the protein-ligand system were constructed in cubic box using PYMOL (61). The systems were then solvated with TIP3P water molecules and neutralized using 150 mM NaCl. All the systems were energy minimized and equilibrated to a temperature of 310 K and 1 atm pressure before proceeding to production runs. Velocity rescaling algorithm (62) with (time constant of 0.1 ps) and Parrinello-Rahman algorithm for (63) (time constant of 2 ps) were used to maintain temperature and pressure respectively. Bond constraints were dealt using LINCS algorithm (64). The long-range electrostatic interactions were dealt using particle mesh Ewald scheme with order 4 and a Fourier spacing of 0.16 while the short-range interactions were dealt using Verlet scheme with a cutoff of 1.4 nm. All the simulations were performed in GROMACS-5.1.4 (65) patched with PLUMED-2.3.5 (66). Short production runs of 10 ns were performed before proceeding for metadynamics runs. The details of different systems and the definition of their respective collective variables can be found in SI as well as in the input files shared on the Github repository.

### Umbrella sampling simulations

The initial co-ordinates for setting up the umbrella sampling simulation were taken from the last frame of the corresponding PHD (WT/mutant)-lipid all-atom trajectory at the end of 500 ns runs. To get the umbrella locations, we placed PHD away from the membrane surface and saved the co-ordinates of the system at every 1 Å window upto a distance of 25 Å . The systems were then ionized and solvated for AAMD runs. The co-ordinates thus obtained were used as initial configurations for the 25 windows that constitute umbrella sampling simulations. Here, we choose the distance between the center of masses of the protein and the membrane to be restrained as the window centres (with restraint constant k = 1000 kJ/mol/nm^2^). Each window was simulated for 50 ns of production run with the umbrella restraint turned on preceded by an equilibration run. The potential of mean force (PMF) profiles were then calculated using GROMACS g_wham tool. The error along the PMF profile was obtained by performing bootstrap analysis with 100 bootstraps. All profiles were shifted so that the PMF value at the membrane plane was at 0 kcal/mol value for reference. The same protocol was used for generating the PMF profiles of all the systems of interest.

## RESULTS AND DISCUSSION

### Binding of the PHD renders the membrane pliable for fission

To determine the effect of the PHD on the mechanical properties of the membrane, we performed 4 *μs* long coarse-grained (CG) simulations of a large lipid bilayer patch (~ 625 *nm*^2^) using the Martini force field (47). This bilayer is composed of 2048 lipids with DOPC:DOPS:PIP2 in 80:19:1 molar ratio. 14 CG PHD units were mounted randomly on to one of the leaflets of the bilayer. The bending moduli calculated by fitting undulation spectra to regular Helfrich theory (see Fig.2) show that the presence of the PHDs reduces the bending modulus of the lipid bilayer. We notice that the bending modulus (*K_c_*) of the membrane with PHDs is about 4.5 *k_B_T* lower than without PHDs. To gain insights into the molecular structures and changes related to the two-dimensional density structure factor of the membrane, we plotted the full height undulation spectra (see Fig. S1 in SI) with larger q-regimes (49). We observed some differences between the two spectra in the intermediate q regime but no noticeable differences in in-plane molecular structure fluctuations in the larger q regime when interpreting and comparing the full spectra. Analyses of trajectories did not reveal any noticeable protrusion modes, which we had anticipated from the differences that we saw in the intermediate q-regimes.

**Figure 2:**
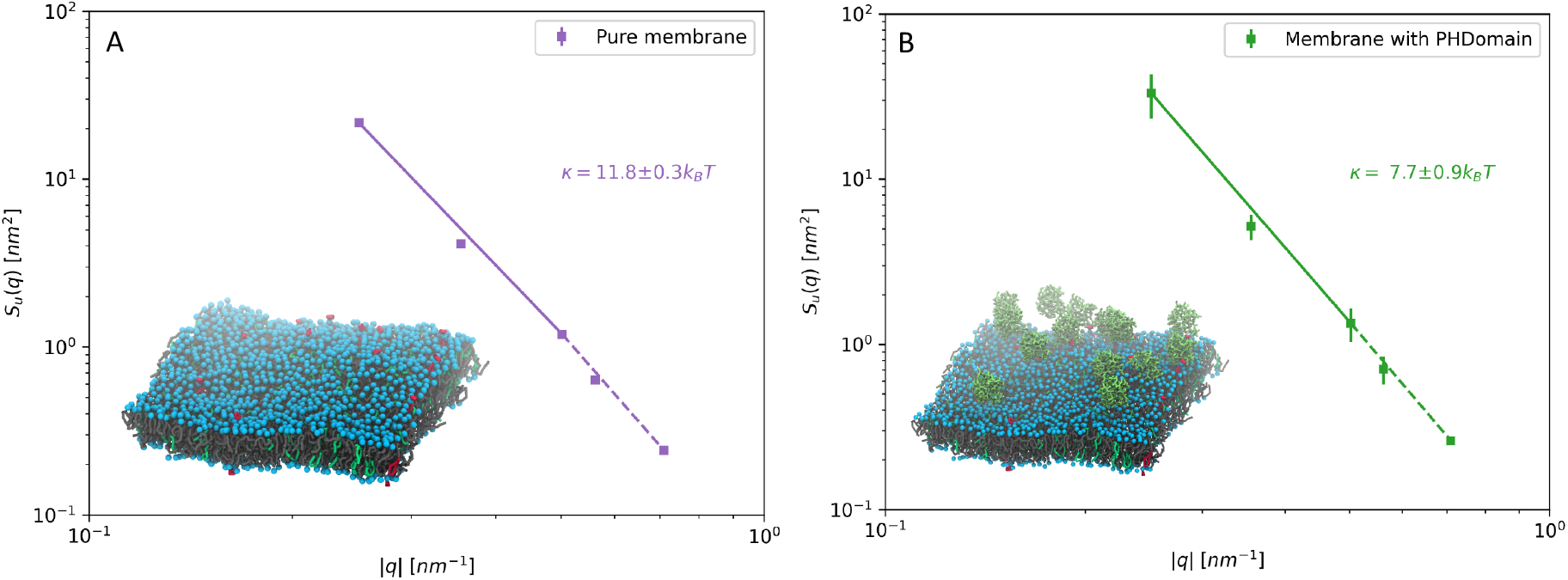
Height-height undulation spectra for the two Martini CG systems (A) Spectra with 2048 lipids and (B) with membrane and 14 CG PHDs distributed randomly on the bilayer surface. The schematics of the system, rendered from snapshot of the trajectory, are shown in inset. The undulation spectra follow the standard Helfrich type *q*^4^ scaling and the effect of curvature on bending modulus is not significant. Full fluctuation spectra for coarse-grained systems comprising of 2048 lipids with pure membrane (no PHD) and membrane with 14 PHD are shown in Fig S1.

Based on our finding that the presence of the PHD lowers the bending modulus of the membrane, we wanted to explore the molecular origin of decrease in bending modulus. For this, we generated a microsecond-long atomistic trajectory of a single PHD on a lipid bilayer composed of DOPC: DOPS: PIP2 (80:19:1) and observed the membrane thickness and lipid conformations (tilt and splay) of lipids proximal and distal to the PHD on the membrane. Fig. 2A shows a snapshot of the bilayer-PHD system with the proximal and distal lipids marked in different colors. We define lipid splay as the distance between the terminal carbon atoms on each tail of the lipid. Lipid tail tilt is defined by the angle between the membrane normal and the vector formed by the glycerol carbon and the last tail atom on the chains. We report the average of the two values per lipid for tail tilt. Lipid head tilt is defined by the angle between P-N vector (vector formed by Phosphorus and Nitrogen atom in the lipid) and the membrane normal. Fig 3B shows the variation of membrane thickness across the cross-section of the bilayer averaged over the last 250 ns-long time period. Remarkably, the PHD influences the underlying membrane by thinning it by as much as 0.4 nm (Fig. 3B), which would have a bearing on the bending modulus (67). Similar analyses for different 250 ns-long time-averaged plots show that membrane thinning is tightly coupled to the location of the PHD (Fig. S2, top panel). The change in thickness comes about due to changed molecular configurations of lipids proximal to the PHD. Indeed, presence of the PHD leads to a positive lipid splay in proximal lipids (68) and is seen by measurements of the lipid tail (Fig. 3C) and head (Fig. 3D) tilt angles for a 25 ns time period. As seen in Figs. 3C and 3D, presence of the PHD induces significant fluctuations in the tail and head tilt angles in lipids proximal to the PHD than in the distal lipids or in lipids in a membrane without the PHD. Snapshots of tilt and splay for a proximal and distal lipid are shown in Fig. S2 and highlight the extent to which the PHD influences local lipid dynamics. The ability of PHD to induce such noticeable changes in tilt fluctuations of lipids strongly suggests the possibility that its engagement with the membrane could prime proximal lipids to attain non-bilayer intermediates and thereby lower the energy barrier for fission (23, 25, 26, 69). Importantly, the changes seen in the tilt values and the higher populations of tilt angles from the average emphasize the need to go beyond the simple Helfrich-like elastic sheet models of membrane, including augmented models such as ones with undulation-curvature coupling (70), to understand energetics of membrane remodeling in terms of finer molecular degrees of freedom such as lipid tilt and splay (26, 71).

**Figure 3:**
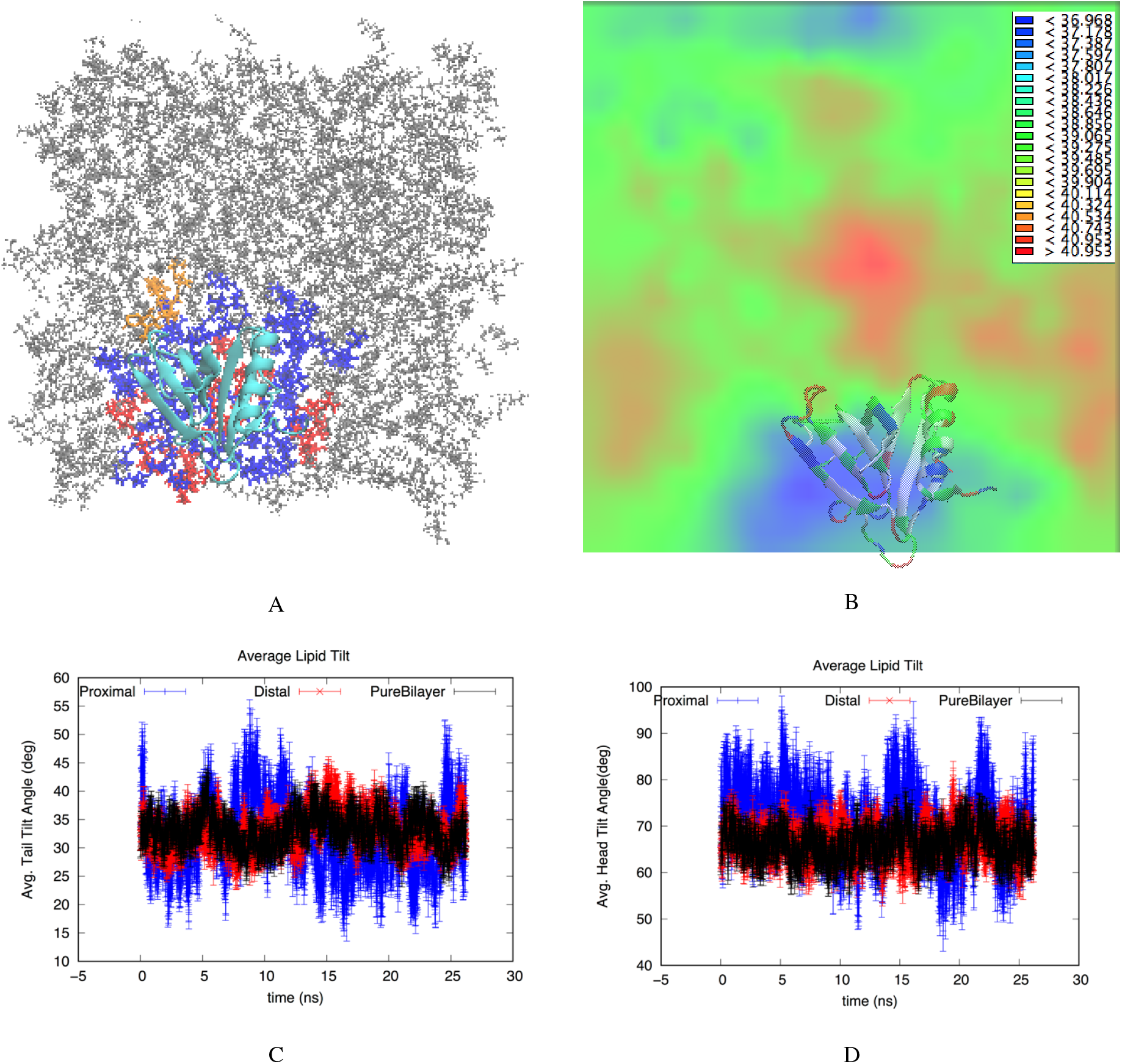
(A) Snapshot from an all-atom DOPC-DOPS-PIP2 bilayer system with single PHD (shown in cyan) on the upper leaflet. The lipids proximal to the protein are marked in blue (DOPC), red (DOPS) and orange PIP2. The distal lipids are marked in grey. The lipids were tracked for 500 ns for the analysis (B) The average thickness profile of the bilayer with the PHD showing thinner regions in blue and thicker regions in red. (C) Average tail angle for lipids proximal to the protein (blue), distal to the protein (red) and in pure bilayer with no proteins (black) (D) Average head angle for lipids. In Fig. S2, we provide some more data with thickness profile of replicates and visuals of lipid conformations near and far away from the PHD.

Together, the abovementioned results confirm that PHDs soften the bilayer and possibly create favorable lipid configurations that induces or stabilizes the non-bilayer intermediate topologies in the course of membrane fission. The implications of these molecular degrees of freedom in terms of the two paradigmatic models of membrane fission, namely the “instability model” (72–74) and the “catalytic model” (23, 26) are discussed in detail later in the text.

### Dynamin PHD engages VL4 loop as an auxiliary pivot that regulates its orientation flexibility

Recent cryoEM reconstructions of the dynamin polymer assembled on a membrane report of a super-constricted state at Å resolution, which highlights localized conformational changes at the BSE and GTPase domains that drive membrane constriction on GTP hydrolysis (27, 28, 75). We reconstructed this data and focused on the PHD orientation by taking slices across the collar, which provided us some interesting observations (Fig. 4A). PHD orientation about a ring of the dynamin polymer was measured by plotting the distribution of the angle between the major inertial axis of each PHD and the vector that connects the center of the ring with the centroid of the PHD (Fig. 4B, see inset in Fig. 4A for a schematic of the angle measurement for a single slice). This analysis revealed that the membrane-bound PHDs adopt a wide range of orientations in the polymer. This is also apparent in other slices analyzed in the polymer (Fig. S3). The wide range of PHD orientations, even in the highly packed constricted state of the dynamin polymer, suggests that the PHD has the inherent ability to associate with the membrane in multiple orientations.

**Figure 4:**
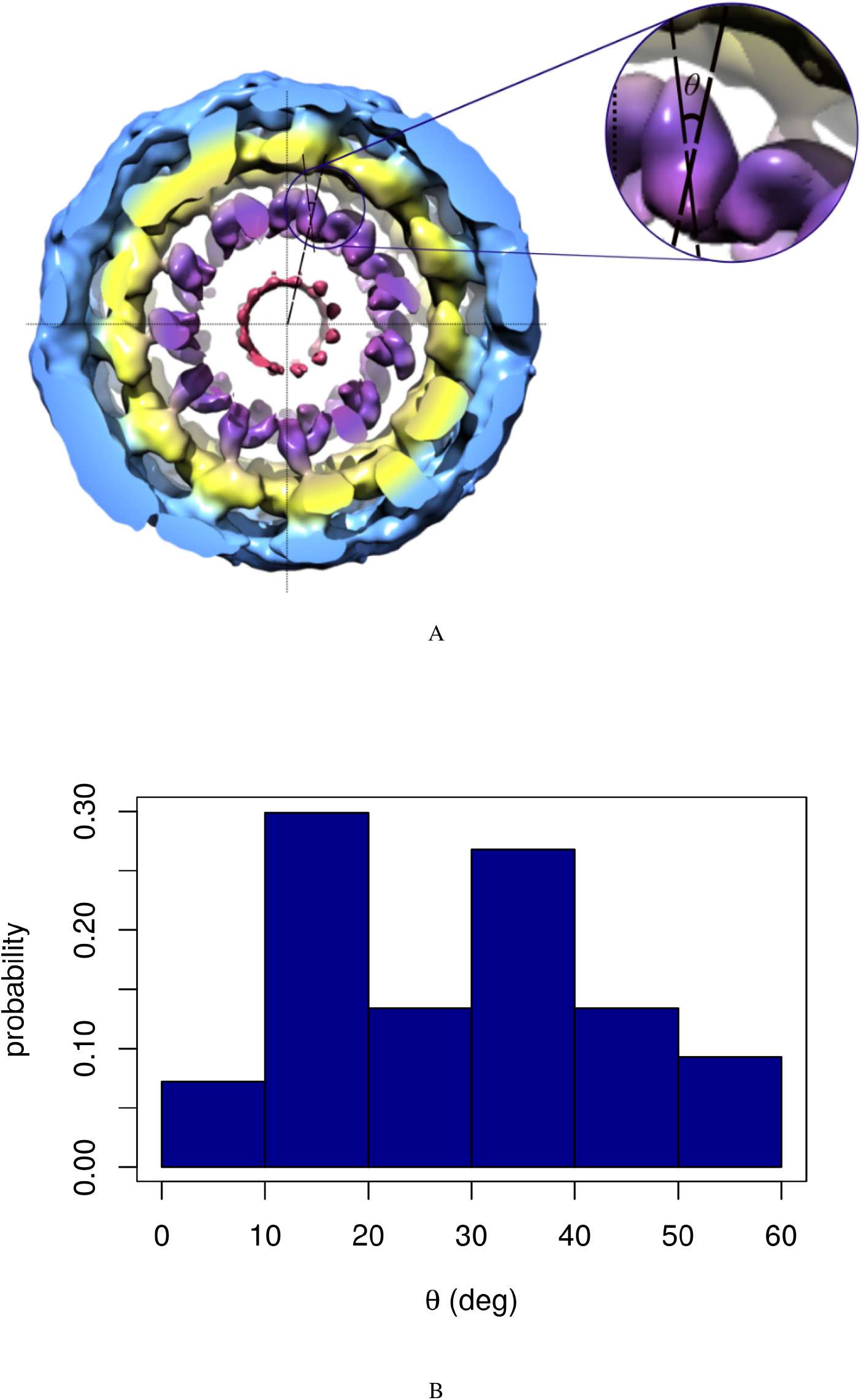
(A) End view of the 3D density map of dynamin polymer assembled on the membrane. The radial densities colored in purple shows the PHDs on the collar. The inset show the angle that the end to end vector of given PHD makes with the radial line passing through the centres of the tube and the given PHD. The density map is re-drawn from the available cryo-EM density data by Jennifer Hinshaw and co-workers (B) The probability distribution of angle described above in the cryo-EM map. The angle *θ* has a wide range for the constricted collar. In Fig. S3, we show reconstruction for six other slices.

To test this and to obtain molecular insights into the association between the PHD and the lipid bilayer, we carried out advanced mixed-resolution molecular simulations using the highly mobile membrane-mimetic model (HMMM) (50, 51, 76). This model utilizes a biphasic setup comprising of lipids with short-chain fatty acyl chains (typically 3-5 carbon atoms-long) organized around an organic solvent, 1,1-dichloro-ethane (DCLE), which mimics the hydrophobic core of the membrane. Many studies have shown remarkable success in probing membrane-protein interactions using HMMM model in MD simulations without compromising on the atomic details of the protein-lipid headgroup interface since lipid diffusion in the bilayer is accelerated by an order of magnitude due to the absence of tail friction (39, 77–79). We ran 12 different HMMM simulations with different initial orientations of the PHD on a bilayer composed of PC:PS:PIP2 (80:19:1 mol%) and a DCLE hydrophobic core. Each system was run for at least 500 ns, with a total simulation time of ~ 9.7 microseconds.

An important observation stands out from these simulations. In all our simulations, we consistently see that PHD associates with the membrane using both (^531^IGIMKGG^537^) and VL4 (^576^EKGFMSSK^583^). VL3 (^590^NTEQRNVYKDY^600^) does not show direct membrane anchoring but is always proximal to PIP2 lipid headgroup that tends to “stick out” of the membrane plane. We explore this feature of VL3 more and put it in context of existing literature (11, 18) when we discuss the binding geometries using the all-atom trajectories. In the figure Fig. 5 below, we have picked the extreme example from our HMMM runs where the initial PHD configuration is set up such that all the loops are facing away from the membrane Fig. 5(a). The final configuration is shown in Fig. 5(b), where the VL1 and VL4 clearly partition into the membrane. Fig. 5(c) shows the time evolution of residues on the various loops and Fig. 5(d) shows the initial and final z-distances for each residue with respect to membrane phosphate plane. We have made a movie file for the full trajectory of this system, which is shared in the SI as MovieHMMM. The VL1, VL3 and VL4 loops are marked as Blue, Red and Orange in color, respectively. VL4 loop shows highly stable membrane association and this observation is consistent across different starting configurations. In Fig. 5(e), we show the convergence data for each of the 12 systems by plotting the distances of the two loops from the membrane surface. Please also see Fig. S4 where the initial and final membrane association profile, in terms of z-distance of each residue away from the membrane plane, is reported for all 12 systems. The initial and final orientation with respect to membrane is also shown in insets of Fig. S4 for each of the 12 systems.

**Figure 5:**
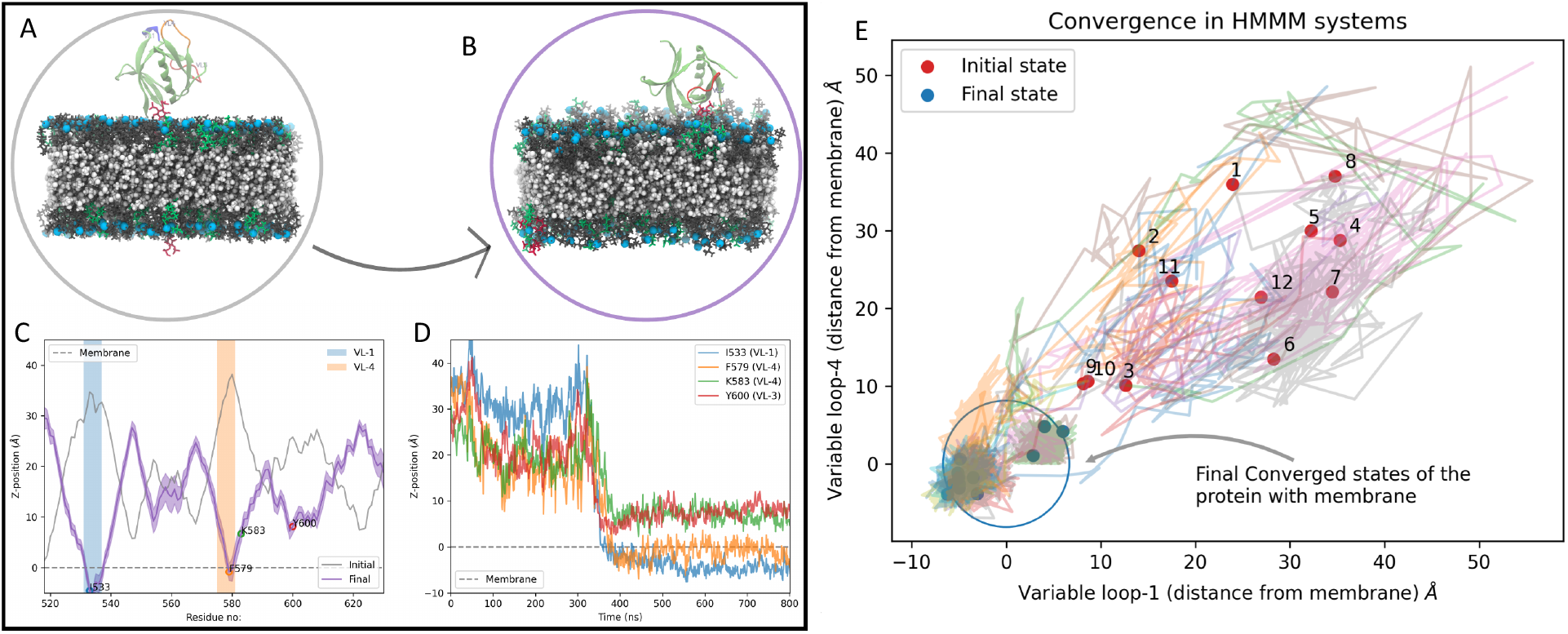
Left panel shows the data for one of the 12 HMMM runs. In this case, the PHD is initially positioned such that all loops face away from membrane. The PC, PS and PIP mimetic lipids are shown in glass grey, green and red, respectively and Phosphorus are marked in light blue. DCLE is shown in white in the core of the bilayer. The VL1 of PHD is shown in blue, VL3 in red and VL4 is shown in orange color. The left panel shows the initial (A) and final (B) PHD configuration. (C) shows the initial and final z-distance of each of the residues averaged over last 100 ns (D) shows the time evolution of signature residues on different loops. The whole 500 ns trajectory data for this system is also reported as a movie file in the SI (MovieHMMM). In the left panel, (E) shows the convergence data for all 12 replicates of the HMMM run, where the distance from the membrane surface for the VL1 loop (I533) on x-axis and VL4 (F579) on y-axis is tracked with time. Almost all system converges to a configuration with VL1 and VL4 physically engaging with the membrane. Plot akin to one on the right panel is reported for all 12 systems in Fig. S4 (i-xii)

Results on VL1 are consistent with previous studies indicating that it acts as membrane anchor and the I533A mutation on VL1 is known to destabilize this interaction and reduce the stability of the scaffold on the membrane in the wake of GTP hydrolysis (3, 14, 15, 17, 31, 44, 80). To get further insights into how the I533A mutant affects the membrane association, we carried out AAMD simulations for WT-PHD and I533A-PHD on a bilayer composed of PC:PS:PIP2 (80:19:1 mol%). Each system was run for at least 500 ns. Fig. 6(a) shows the residue-wise z-distance for initial and final configurations for the WT system. Fig. 6(b) shows the same data for I533A system. To our surprise, the membrane association profile for the I533A did not show noticeable changes and the mutant does not seem to be membrane defective. We also carried out similar AAMD runs on mutations on VL4 residues such as F579A and K583A, which also showed association behavior similar to WT. We anticipated these to be facile association and carried out umbrella sampling calculations on the WT and various mutant systems to extract the binding free energy (Δ*G*) information. Fig. 6(e) shows the membrane dissociation “potential of mean force” (PMF) profile for the WT and various mutant systems. As anticipated and reassuringly, there is a significant difference in membrane binding free energies for the WT and I533A mutant system. This suggests that while the mutant I533A is non-defective with respect to membrane association, it binds superficially to the membrane and that could explain the compromised assembly and reduced stability of the dynamin on the membrane.

**Figure 6:**
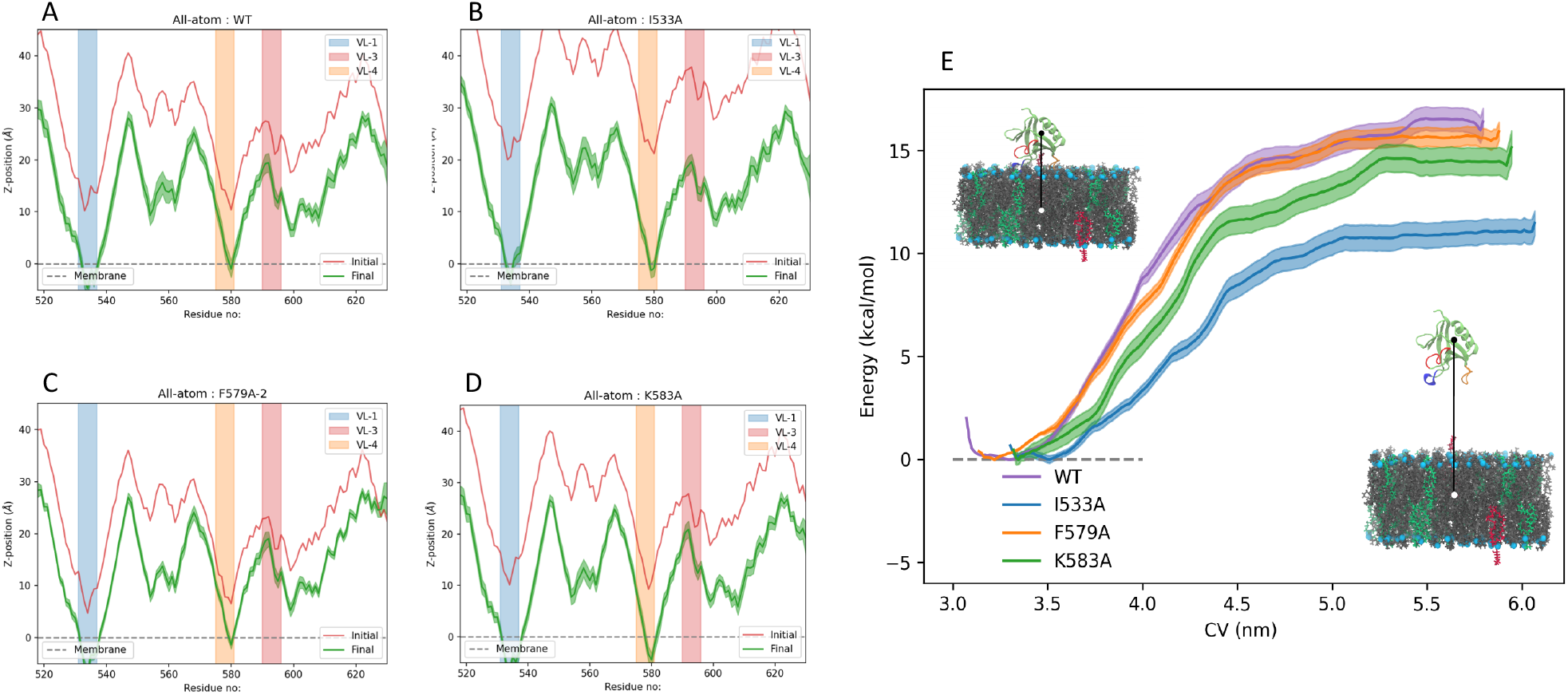
Left panel shows the data for four of the AAMD simulations runs with respect to z-distance of each residue on the protein away from the membrane phosphate plane. The simulations were run for 500 ns each. The four systems are (A) WT, (B) I533A, (C) F579A and (D) K583A. The red curve on each figure indicates the initial configuration and the green curve is the final configuration averaged over last 100 ns. Similar plots for K339A and Y600L mutant systems are shown in Fig. S5. Right panel (E) shows the PMF profiles of the four systems obtained through umbrella simulations runs. Right most end of the graph denoted PHD in solution. The zero of the free energy difference plot is set at phosphate plane of the membrane.

The analyses on F579A and K583A mutations shed some light on the role of VL4 as an auxiliary/secondary pivot. We chose to test the F579A mutation first since our HMMM simulations as well as our AAMD simulations consistently show that F579 has the deepest penetration for VL4. Interestingly, F579A hardly shows any change in binding free energies from the calculations done using umbrella sampling. K583A shows a minimal change of about 2 kcal/mol in the dissociation Δ*G* values as compared to WT. It is clear that mutation on VL4 does not significantly alter the dissociation Δ*G* so we checked if these mutations had any effect on angular stability of the membrane bound PHDs. To quantify the orientation flexibility of membrane bound PHD, we looked at the angle that the PHD helix makes with the membrane normal (*θ*) and also the angle that it makes when projected to a reference axis on the membrane surface (*ϕ*), represented in 360 degrees cylindrical coordinate system. Inset schematics in Fig. 7(a) shows the angles. We compare the angles for WT data against I533A, F579A and K583A (*θ* in Fig. 7(a) and *ϕ* in Fig. 7(b)). The distribution in WT is consistent with our cryoEM reconstructions analysis (27, 28) and further show that that membrane-bound PHD is flexible in its orientation. However, VL4 mutation such as F579A and K583A shows significant increase in the range of angle distribution, while I533A hardly shows any change in the distributions. Possibly, mutation that keeps the PHD anchored but makes it highly labile (orientationally) adversely affects high-fidelity collar assembly leading to compromised fission behavior. So even if, on the face value, the PHD is not membrane defective with VL4 mutants, the high orientation fluctuations is likely to affect the fission kinetics to a large extent. By attaching to the membrane using VL1 as the primary pivot and VL4 loop as the auxiliary pivot, the PHD stabilizes the dynamin on the membrane - both in terms of anchoring and orientation flexibility - thereby likely facilitating better assembly and collar formation. Incidentally, artificially higher PIP2 concentration on the membrane has similar effect on rescuing the kinetics for I533A mutation to some extent (44), which lends further credence to our theory. While a high PIP2 concentration rescues the fission kinetics by increasing the “anchoring” binding energy, it will be interesting to check if fission kinetics can be rescued for mutations that makes the PHD unusually labile on the membrane. We carried out some tests towards that with double mutations as shown in Fig. S5.

**Figure 7:**
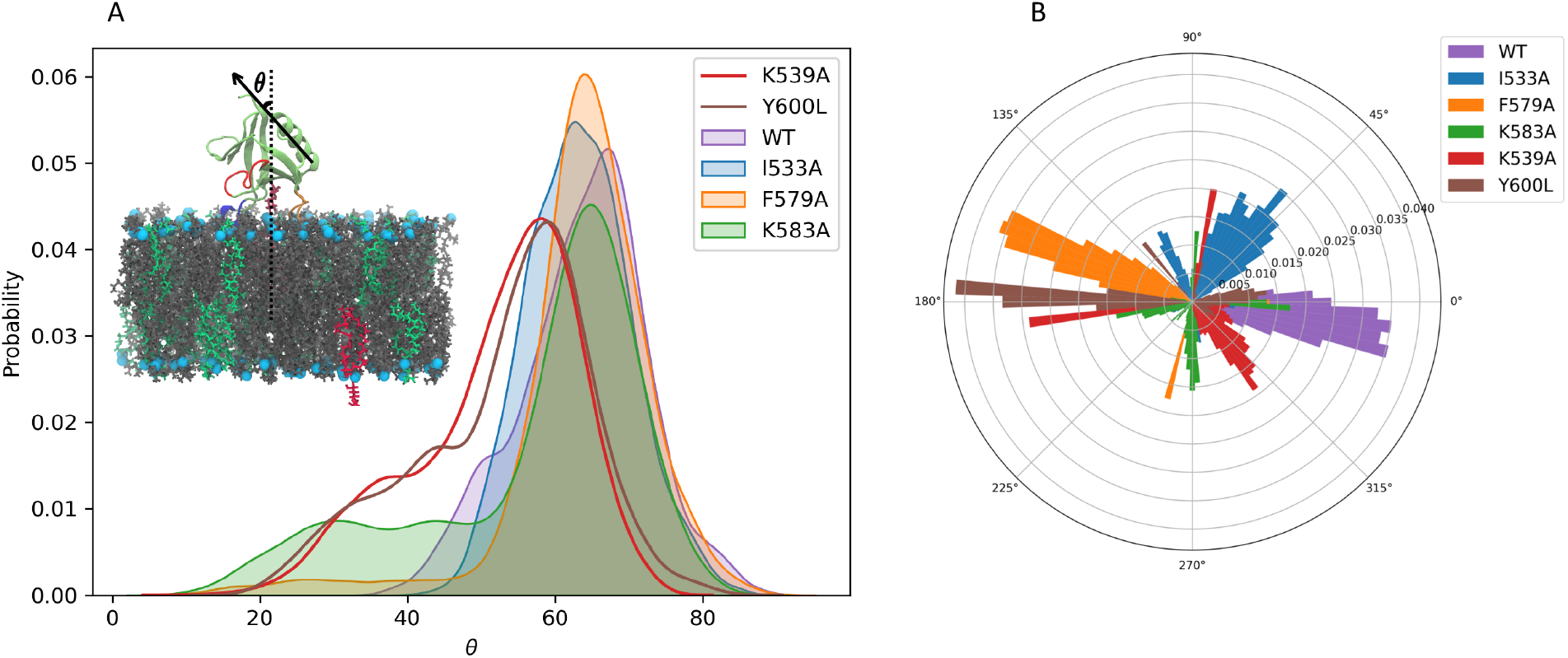
Data showing orientation distribution for PHD for WT and various mutant systems. Inset in the left panel shows a schematic where *θ* is the angle that helical axis makes with the membrane normal. *θ* angle distribution for WT, I533A, F579A, K583A, Y600L and K539A systems are shown in left panel. *ϕ* is the angle that the helical axis makes with a reference axis on the plane of the membrane and is shown the right panel (B) for all the WT and mutant systems.

### Key residues stabilizing the PIP2-PHD binding are found in multiple VLs

Role of VL1 in dynamin-catalyzed vesicle scission is undisputed (3, 17, 18, 23, 44). Fission is shown to be impaired with VL1 mutants such as (I533A, M534C) that attenuate its hydrophobic character (17, 81). Several studies have also shown the importance of VL3 in stabilizing the membrane interaction of PHD (11, 16, 18, 43, 43, 82). Due to its polar nature, VL3 direct engagement with the hydrophobic core is thought to be unlikely and loss in function with mutations such as Y600L is primarily attributed to the overall instability of the dynamin polymer on the membrane surface (18). Additionally, our work consistently shows that VL4 engages with the acyl-chain hydrocarbon core and likely acts as a secondary pivot or a “buoy” that stabilizes the protein on the surface. Cryo-EM maps of the dynamin polymer, reconstructed with different modeled orientations of the PHD further reinforces the possibility of PHD associating with multiple orientation with the membrane (27, 28, 82).

To test the multiple orientation and binding geometries and also to look more carefully into the molecular-scale features that possibly drive and stabilize these association, we carried out well-tempered metadynamics (WT-MTD) advanced-sampling simulations (83–86) using a single PHD and head group mimic of the PI(4,5)P2 (IP3: Inositol triphosphate). We set up three different systems for WT-MTD simulations: two systems where the distance collective variables (CVs) are chosen such that the variable loops are sampled preferentially and another system with no bias for any loop or pocket. CV can be thought of as reaction coordinates or order parameters that can distinguish between different conformational states of the PIP2-PHD complex in this work. The schematics of CV selection is shown in left panel of Fig. 8 and the corresponding free energy surface (FES) profiles are shown in the right panel. The details of CV definition is discussed in the SI. The convergence plot for the three systems and the time evolution of distance in and out of the ligand from domain’s pocket is show in Fig. S6. In order to explore if a variety of binding geometries exist, we considered the geometries in the range of 2 *k_B_T* of the deepest minima. We observe a range of possible geometries suggesting degeneracy, which implies that *PIP*_2_, or rather *IP*_3_ ligand, can bind with different sets of residues in the PHD. Results from IP3-PHD simulations show that pockets formed by VL1-VL3 and VL1-VL4 are both available for binding the inositol headgroup. It should be noted that the WT-MTD runs were carried out with isolated lipid headgroup (IP3) and so we wanted to check how well the information from these runs would relate back to membrane interaction modes of the PHD. To address this issue as well as to gain further insights into the configurations of binding geometry of bilayer bound PHD, we performed the following simulations. We extracted the co-ordinates of PHD-IP3 from the deepest basins of free energy landscapes obtained from the WT-MTD runs. First set of coordinates were obtained from WT-MTD data where the *IP*_3_ was positioned in the pocket between the VL1 and VL4 loops. The other set of coordinates were taken from simulations where the *IP*_3_ was positioned in the pocket between the VL1 and VL3 loops. The coordinates were superimposed onto the membrane bilayer (PC:PS:PIP2 in 80:19:1 ratio) making sure that the orientation of PHD-*IP*_3_ remains unchanged. This was done by aligning the *IP*_3_ ligand with the head group of PIP_2_ the membrane with only rotational and translational modes allowed during superimposition. The prescription is similar to what was done with EPR-based structure of IP3-PHD of GRP1 to simulated a bilayer bound PHD (32, 87). We then let the system evolve in an all-atom NPT ensemble simulation for 500 ns each. This allowed us to test the stability of the configurations obtained from IP3-PHD WT-MTD runs in the presence of bilayer. We find that the simulation initiated from the orientation taken from minima obtained in the region of loops 1-3 is unstable compared to the orientation observed in the minima obtained from loop 1-4 system. Please see Fig. 9(a) where RMSD is plotted keeping the first frame (configuration from converged WT-MTD runs) as the reference indicating that VL1-VL4 basin configuration is robust. We chose this system for further analyses to look into residues that consistently engages with the acyl-chain hydrocarbon core as well as residues that have non-transient interactions with *PIP*_2_ lipid as well as explore if any of the interactions localizes the anionic PS lipids and stabilizes the interactions further.

**Figure 8:**
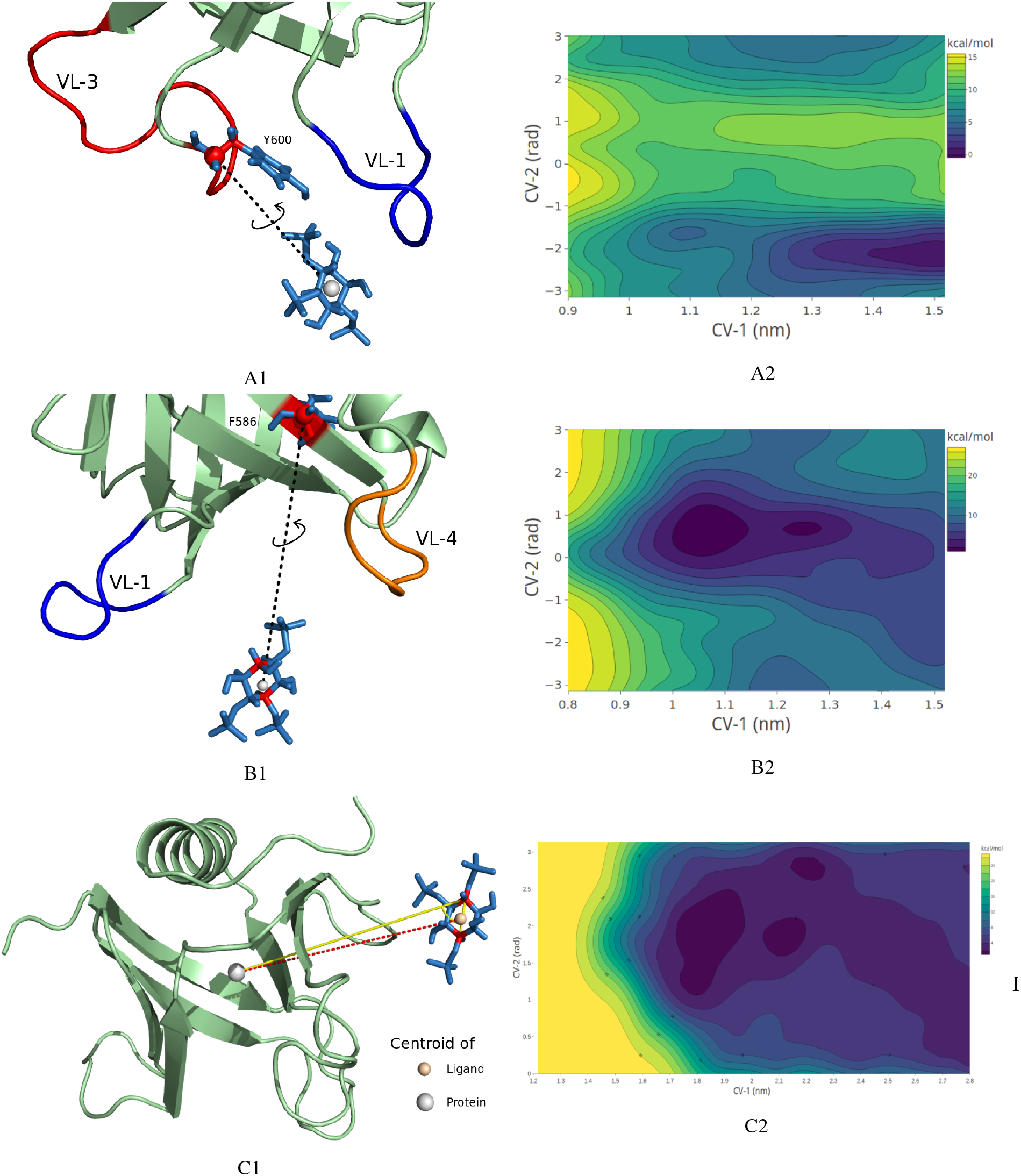
Well-tempered metadynamics setup with docking region for IP3 (ligand) biased towards VL1 and VL3 loop region (A1), towards VL1 and VL4 loop region (B1) and with no bias towards any loop region (C1). The free energy landscape for the three systems are shown in panels A2, B2 and C2, respectively. The lowest energy configurations are picked from the lowest basins and regions free energy difference of < 2*k_B_T* from the minimum basin.

**Figure 9:**
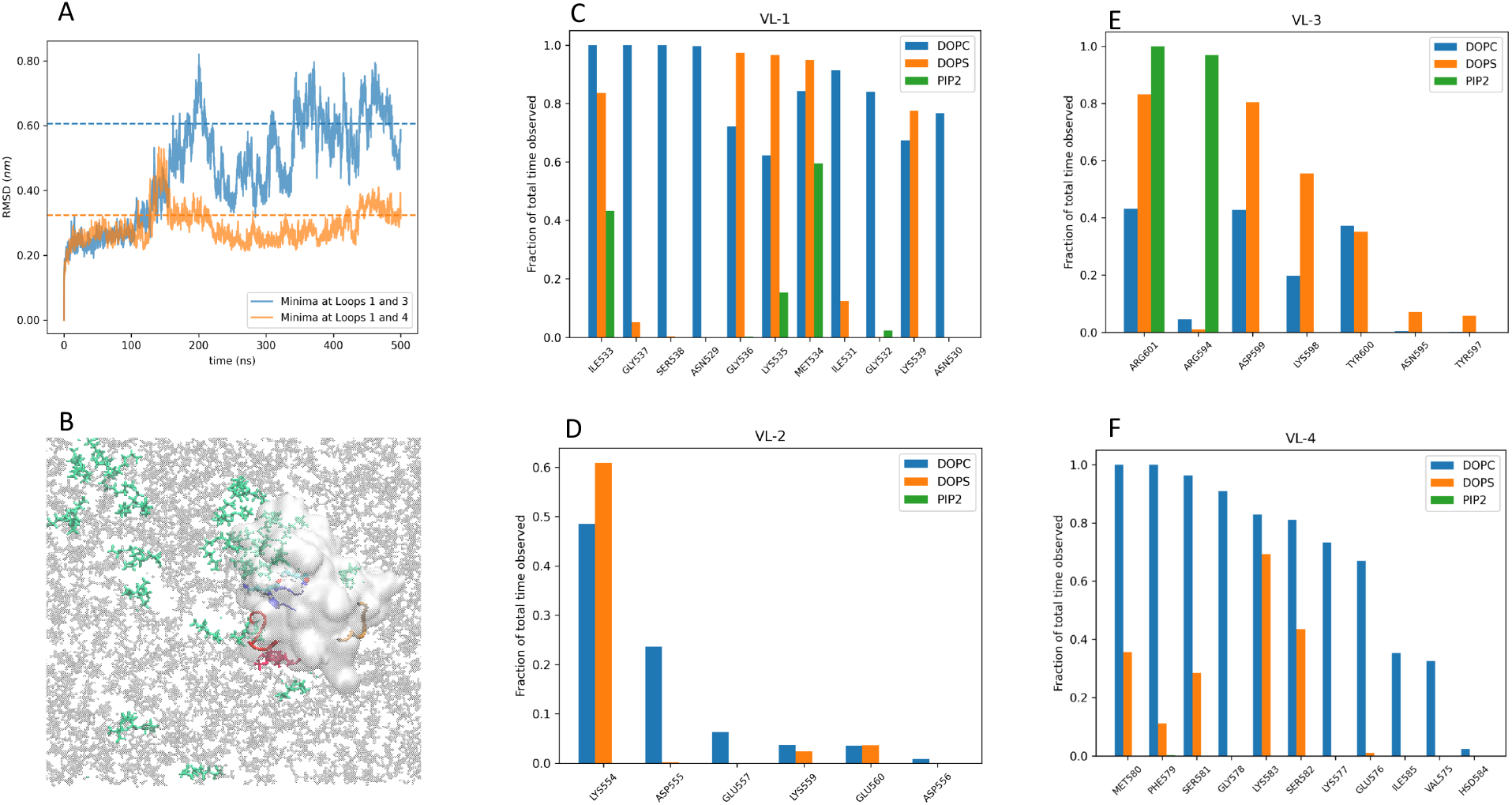
(A) RMSD plot of AAMD runs for two systems, one started with PIP2 docking derived from metadynamics run with IP3 ligand in VL1-VL3 and the other with IP3 ligand in VL1-VL4 pocket. The later shows stable convergence and was chosen for 500 ns of MD simulation, where we explored the lipid contacts of the PHD. (B) shows one of the last snaphots of the MD run (water and ions not shown). PC lipids are shown in Grey, PS in Green and PIP2 lipid in Red. The PHD is shown in a transparent white Isosurf representation and VL1, VL3 and VL4 are shown in opaque Blue, Red and Orange, respectively. The full movie file of this trajectory is also reported as movie file MoviePS. Panel on the right (C-F) shows the normalized contacts that different kind of lipids make with residues on various loops

In Fig. 9(b), we show a snapshot at the end of the 500 ns, which shows the lipid engagements of various loops. Contact analysis of our trajectories show that lipid tail atoms consistently interacts with the residues I533 and M534 of VL-1 (9(c)). That VL1 loop associates hydrophobically is not novel, however what we find very interesting is how the PHD is stabilized further using residues such as K535 and K539 that recruit PS lipid near the VL1. This is quite evident in the movie file MoviePS, which we provide in the SI. VL1 accesses the hydrophobic core in a non-specific manner but once the PS lipids and *PIP*_2_ is recruited to the location, the I533 and M534 residue stabilize PS and *PIP*_2_ in its place by contact with its tail. It is also amply clear from our HMMM simulations (see MovieHMMM) and AAMD simulations (see MoviePS) that VL3 have vital interactions with *PIP*_2_. HMMM runs show that the *PIP*2 is locked to the binding pocket once it makes contact with the polar VL3. AAMD analyses also highlights how *PIP*_2_ is localized without VL3 partitioning into the membrane. A plot of the z-distance of the C-alpha and z-distance of the terminal ‘hydroxyl oxygen’ atom of Y600 as a function of time reveals nominal association with the membrane. While Y600L mutation is tried and tested for fission inhibition behaviour, our data also show that R594 and R601 as critical residues on VL3 (Fig. 9(d)) for *PIP*_2_ interactions and could be tested experimentally to provide further insights into dynamin’s membrane association mechanisms. While VL1 and VL3 loop has been studied in depth for over a decade now (and VL2 is not important for membrane association but shows some role in PS recruitment (Fig. 9(e)) through K554), no study has so far looked into the role of VL4. From our analyses, we show that VL4 consistently partitions into the hydrophobic core of the membrane though the association seems facile given that we see only marginal loss in binding free energies with mutations on VL4 residues. However, mutations such as F579A and K583A have very noticeable effects on the orientation stability of the PHD on the membrane despite no visible loss in membrane partitioning. Fig. 9(d) clearly shows the heavy contacts that F579 and M580 and other residues of VL4 make with the membrane. Our observations can be used inform experiments to test the role played by VL4 in terms of orientation stability of PHD.

## CONCLUSIONS

The dynamin collar undergoes a series of segmental rearrangements while executing membrane fission. Each PHD must be capable of adapting to the progressively remodeled underlying membrane. Previous results from 3D reconstructions of dynamin mutants trapped in a constricted state (27, 28, 82) as well as data from biochemical and multiple fluorescence spectroscopic approaches (18) suggest that the PHD can associate with the membrane in different orientations. Theoretical analyses of determinants that could lower the energy barrier for fission have invoked tilting of the PHD to conform with the evolving membrane curvature and thereby create a low-energy pathway for fission (23, 29, 88). Our multiscale simulations data validate these possibilities and provide molecular insights in terms of effects of PHD on membrane properties and lipid conformations and as well on the binding geometries and orientation stability of PHD on membrane, all of which may affect processes downstream of membrane binding.

We show that binding of the PHDs lowers the membrane bending rigidity indicating that molecular interactions at the membrane interface confer a change in the bending rigidity of the membrane. These changes are brought about by polar and charged residues at the membrane interface that locally reduce membrane thickness and increase chain flexibility. Recent evidence of the existence of non-bilayer configurations of lipids formed at the stalk and hemi-fission intermediates prompted us to go beyond models that assume the membrane as an elastic sheet capable of undergoing fission through curvature instability (11, 72, 74, 89–91) and focus on the molecular degrees of freedom of constituent lipids. Thus, higher fluctuations in the lipid tilt and lipid tails splay may have a significant role in facilitating the membrane to stochastically crossover to the non-bilayer intermediates once a threshold constriction of the lumen is reached. The formation of non-bilayer intermediates dictates that a physical coupling between the two leaflets of the bilayer is reduced (92). However, our analysis reveals no apparent change in the interdigitation between the two leaflets and this could reflect the fact that our simulations are currently not designed to recapitulate dynamics of the constricted tubular intermediate seen during fission.

The PHD is a ubiquitous membrane-binding domain for many different proteins. Several studies have explored the molecular mechanism by which it associates with PIP_2_/PIP_3_ in the membrane (32–41). Analysis of inositol-bound PHD structures clearly reveal the presence of a highly basic region that favors binding to the highly anionic phosphoinositide lipids (30, 93). Our results from AAMD, WT-MTD and HMMM runs extend these models and provide molecular-level insights into dyn-PHD engagement with membrane. Our simulations studies show that membrane binding of dyn-PHD has features that are quite unique. In general, multiple binding sites studies suggest mechanism with canonical and atypical phosphoinositide-binding sites that allows efficient switching between active and inactive forms in proteins (33, 40–42). There is an important difference between the degeneracy in binding sites in our work as compared to that observed in other systems including ASAP1-PHD. Residues from multiple variable loops engage with the *same* PIP_2_ lipid in case of dyn-PHD. The “catalytic” role of dyn-PHD role is more “mechanical” in nature and the multiple binding loops around a PIP_2_ may explain how the dyn-PHD is able to “dynamically” keep itself anchored to the membrane while undergoing very rapid shape changes in the midst of fission process.

Despite the fact that all three VLs appear to engage with the membrane, albeit to different extents and through different modes, the degeneracy in orientations lends credence to the notion that unlike being rigidly clamped to the membrane, the PHD are best represented as units that are flexible and “moored” to the membrane. VL1 acts as a primary anchor that dictates the binding affinity with the membrane, a feature driven by both hydrophobic interactions (I533A showing reduced dissociation free energy) as well as stabilization by recruitment of PS lipids at VL1 by residues such as K535 and K539 on the loop. Interestingly, residue K554 on VL2 seems to also favor PS lipid recruitment to the PHD though VL2 is quite distant from the membrane surface. VL3, a highly polar loop, never partitions with the bilayer but is instrumental in localizing PIP_2_. Given that the protruded headgroup of PIP_2_ allows such interactions, it would be interesting to see how mutations on VL3 residues behave in membrane with different curvatures and anionic lipid composition. VL4 has a subtler role in dictating the orientation flexibility of dyn-PHD and we provide some testable hypothesis towards that end. None of the experimental studies have probed the role of VL4 in PHD-membrane interaction possibly since the center of the loop is rich in hydrophobic residues with distal Lysines unlike VL1 and VL3 (31) where the positive charged residues are at the tip of the loop. Our study identifies this novel interaction between VL4 and the membrane and highlights its role as an auxiliary pivot providing orientation stability to the PHD - a feature we believe is important for effective polymerization of dynamin collar. Our findings with respect to both multiple binding sites (40–42) and multiple orientations (18, 23) have to be relooked into the light of the auxiliary pivot theory. Put together, our results indicate the membrane binding and orientation stability is tightly regulated by a concerted interactions between three different loops on the dyn-PHD, which acts as flexible pivot and modulate effective and expedited fission.

## Supporting information

HMMM movie file

## AUTHOR CONTRIBUTIONS

AS conceived the idea and designed the experiments. KJ performed and analysed the HMMM simulations and the US simulations. KJ and KB processed the recently published cryo-EM electron density map for dynamin collar on the membrane, reconstructed and analysed the images. KB carried out the plain (not reported) and well-tempered metadynamics simulations, multiple all-atom simulations with WT and mutant systems, analysed the data and was the primary person for making the plots for all analyses. AS setup and ran the large scale CGMD Martini simulations, which was analysed by KB. AS wrote the paper with help of all co-authors.

## ACKNOWLEDGMENTS

AS would like to thank Dr. Ryan Bradley from Prof. Ravi Radhakrishnan’s lab at University of Pennsylvania for his help in the analyses and interpretations of the membrane fluctuation spectra. KJ and AS would like to thank Shashank Pant for his help in setting up HMMM simulations. AS would like to thank Vikas Dubey, a junior research trainee in the laboratory, for help with CGMD simulation runs. VD was supported partially by grant from a J. C. BOSE project (DSTO/RV-2052). AS would also like to thank Prof. Raghavan Varadarajan and Prof. Thomas Pucadyil for carefully reading the manuscript and for their comments. Financial support from the Indian Institute of Science-Bangalore and the high-performance computing facility “Arjun” setup from grants by a partnership between the Department of Biotechnology of India and the Indian Institute of Science (IISc-DBT partnership programme) are greatly acknowledged. A.S. thanks the startup grant provided by the Ministry of Human Resource Development of India and the early career grant from the Department of Science and Technology of India. FIST program sponsored by the Department of Science and Technology and UGC, Centre for Advanced Studies and Ministry of Human Resource Development, India is gratefully acknowledged by the authors.

## SUPPLEMENTARY MATERIAL

An online supplement to this article can be found by visiting BJ Online at http://www.biophysj.org.

