## Supporting Material PDF file for "Flexible pivoting of dynamin PH-domain catalyzes fission: Insights into molecular degrees of freedom"

K. K. Baratam[1†], K. Jha[1†], , and A. Srivastava[1*]

[1]Institution A, Address A

Running title: *Dynamics of the membrane-bound dynamin PHD*

† Contributed equally

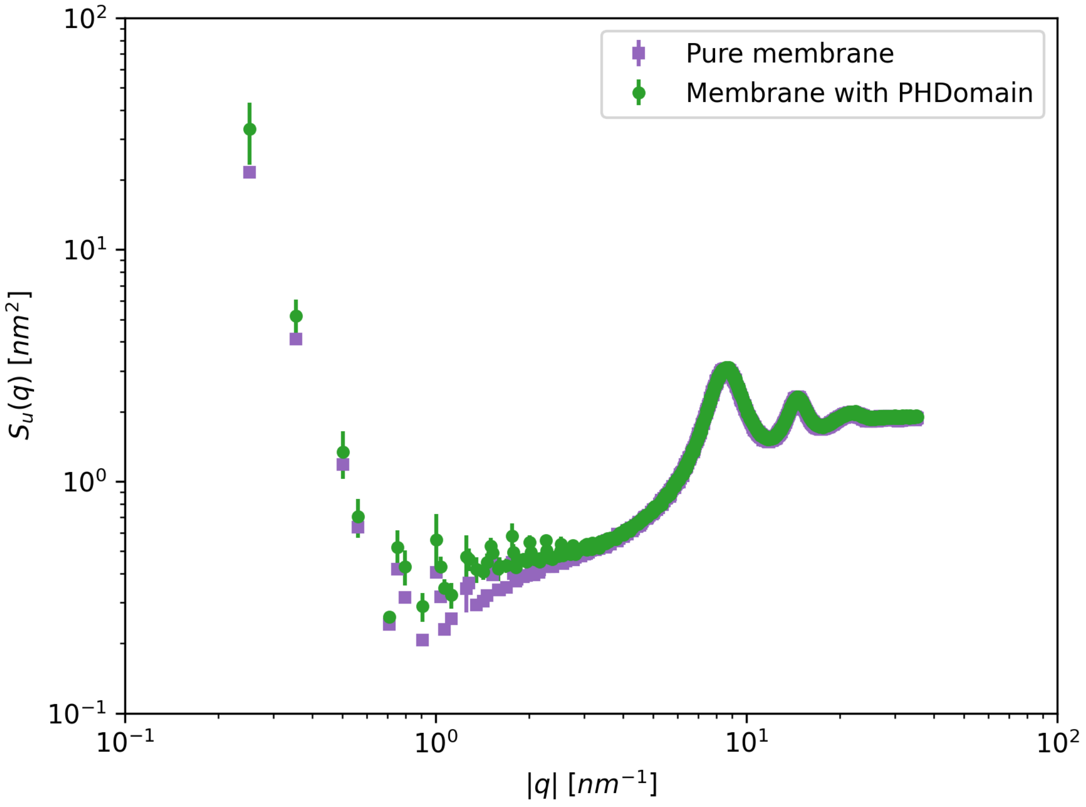


**Figure S1:** Undulation spectra of CG bilayer (2048 lipids Martini forcefield). Pure bilayer spectra is shown in purple while the spectra with 14 PHDs on the top leaflet (randomly arranged) is shown in green.

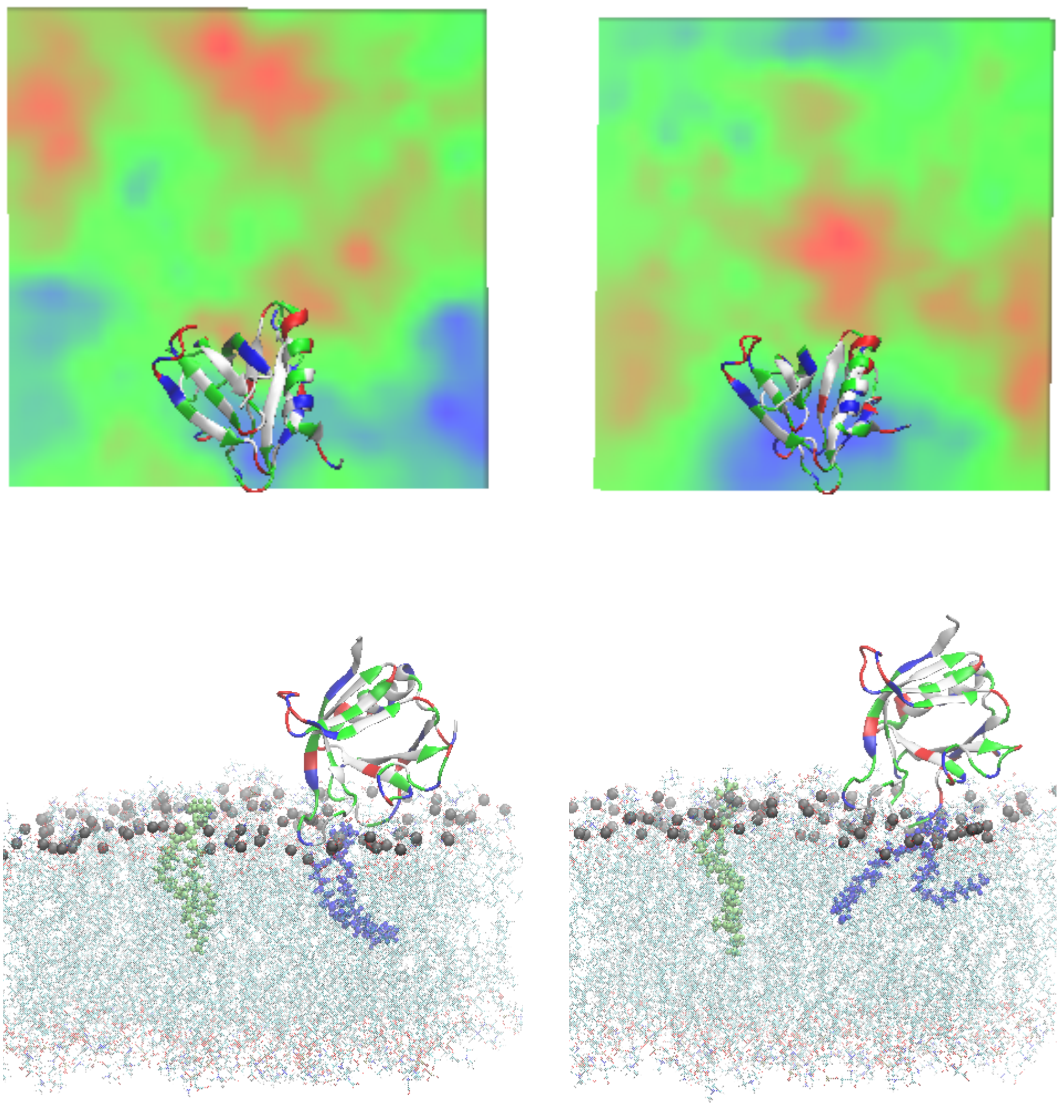

**Figure S2**: Top panel shows the membrane thickness profile for two other systems with dyn-PHD on all-atom bilayer. The difference between the thinnest and thickest regions is around 0.4 nm. The bottom panel shows snapshot of a PHD-bilayer run and the blue lipid denotes a dyn-PHD proximal lipid while the green color shows the distal lipid. The proximal lipid has a visible tilt (left bottom) and very noticeable splay. The average statistical data for the splay and tilt are shown in Fig. 2.

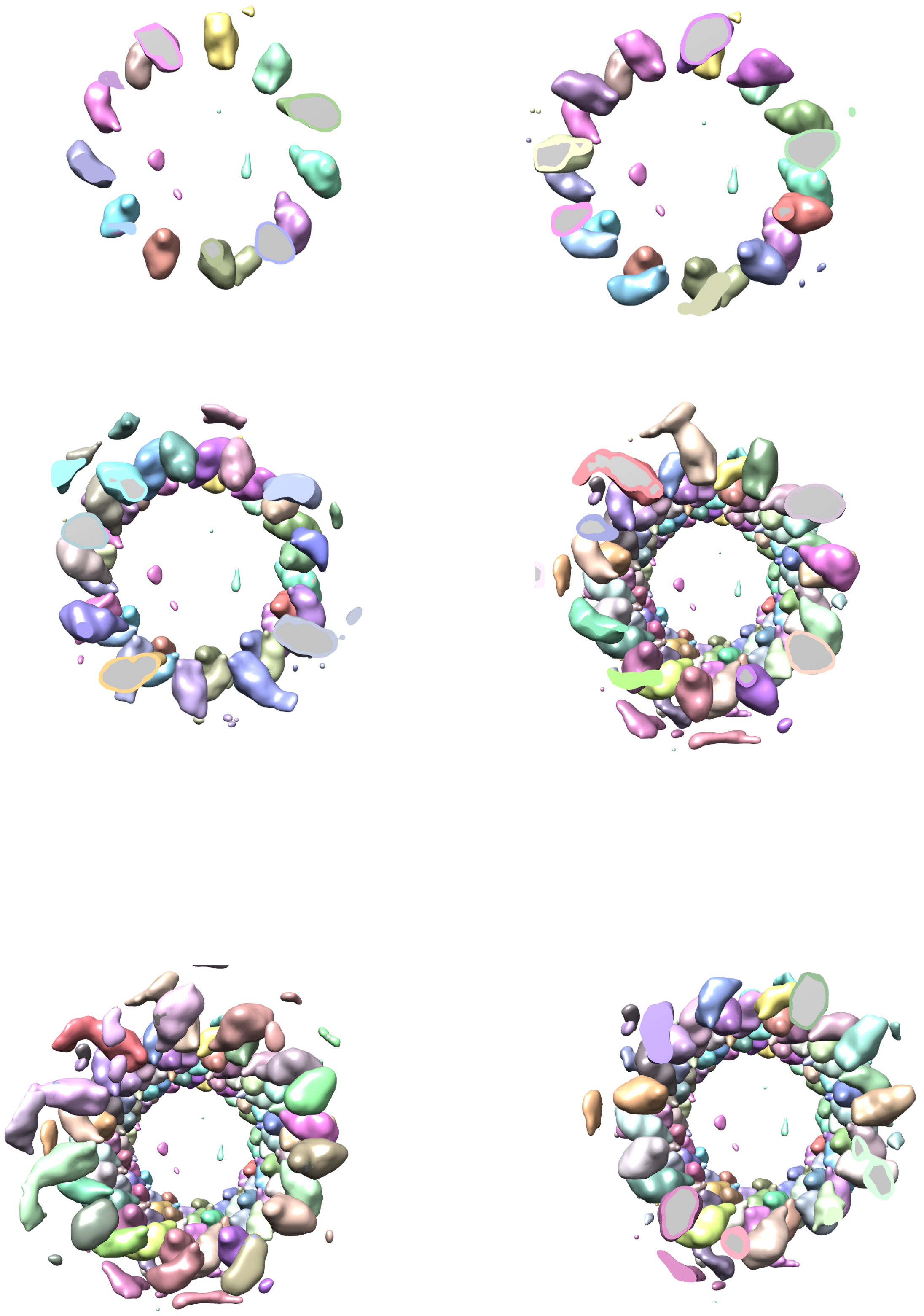

**Figure S3:** End view of the 3D density map of dynamin polymer assembled on the membrane. The top-left panel shows the front most slice of the scaffold with dyn-PHDs in different orientations. The right-bottom panel shows the full scaffold. The density map is re-drawn from the available cryo-EM density data by Jennifer Hinshaw and co-workers

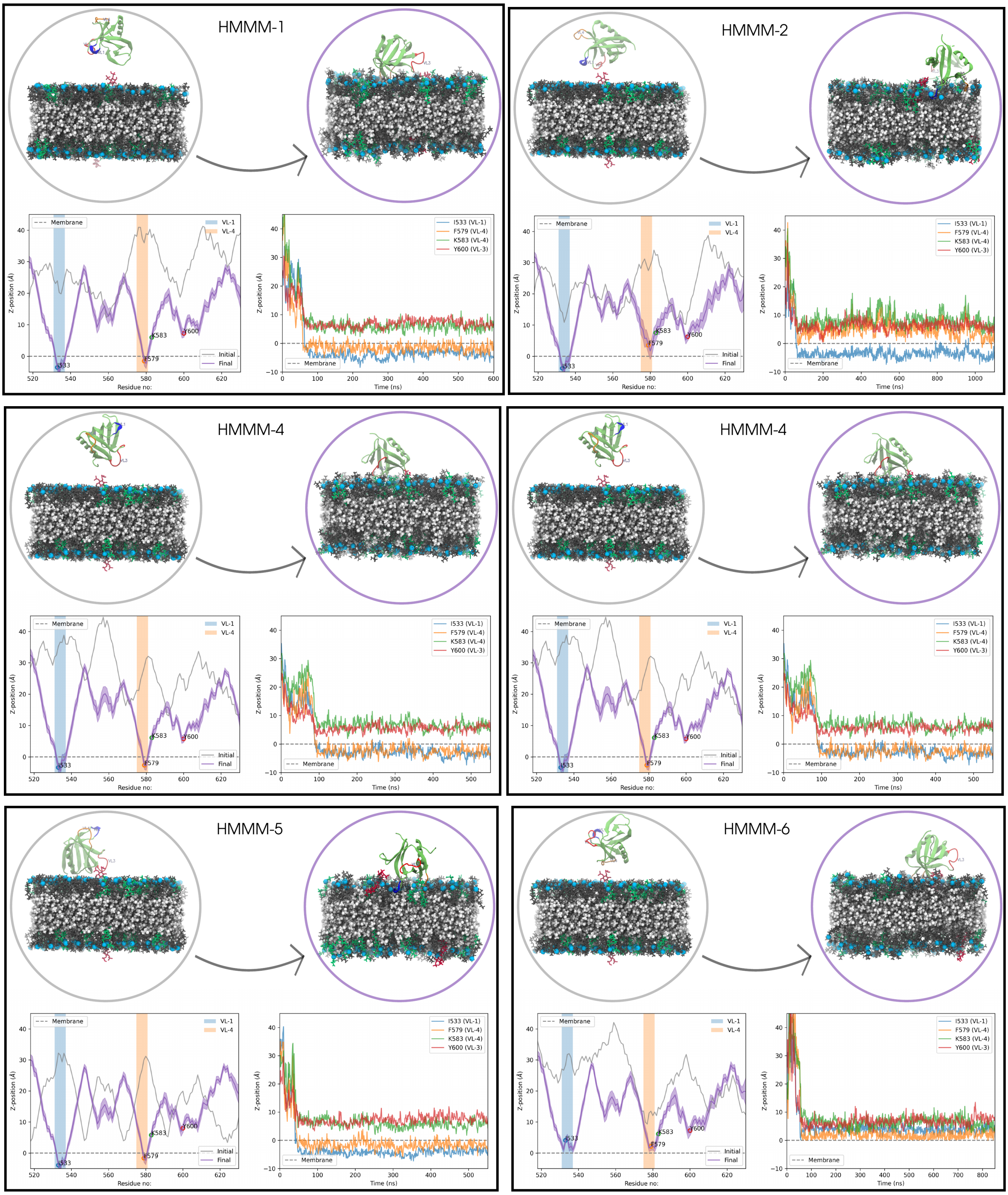
HMMM-1
Membrane
VL-1
VL-4
Z-position (Å)
I533
F579
K583
Y600
Initial
Final
Residue no:
I533 (VL-1)
F579 (VL-4)
K583 (VL-4)
Y600 (VL-3)
Time (ns)
HMMM-2
Membrane
VL-1
VL-4
Z-position (Å)
I533
F579
K583
Y600
Initial
Final
Residue no:
I533 (VL-1)
F579 (VL-4)
K583 (VL-4)
Y600 (VL-3)
Time (ns)
HMMM-4
Membrane
VL-1
VL-4
Z-position (Å)
I533
F579
K583
Y600
Initial
Final
Residue no:
I533 (VL-1)
F579 (VL-4)
K583 (VL-4)
Y600 (VL-3)
Time (ns)
HMMM-4
Membrane
VL-1
VL-4
Z-position (Å)
I533
F579
K583
Y600
Initial
Final
Residue no:
I533 (VL-1)
F579 (VL-4)
K583 (VL-4)
Y600 (VL-3)
Time (ns)
HMMM-5
Membrane
VL-1
VL-4
Z-position (Å)
I533
F579
K583
Y600
Initial
Final
Residue no:
I533 (VL-1)
F579 (VL-4)
K583 (VL-4)
Y600 (VL-3)
Time (ns)
HMMM-6
Membrane
VL-1
VL-4
Z-position (Å)
I533
F579
K583
Y600
Initial
Final
Residue no:
I533 (VL-1)
F579 (VL-4)
K583 (VL-4)
Y600 (VL-3)
Time (ns)

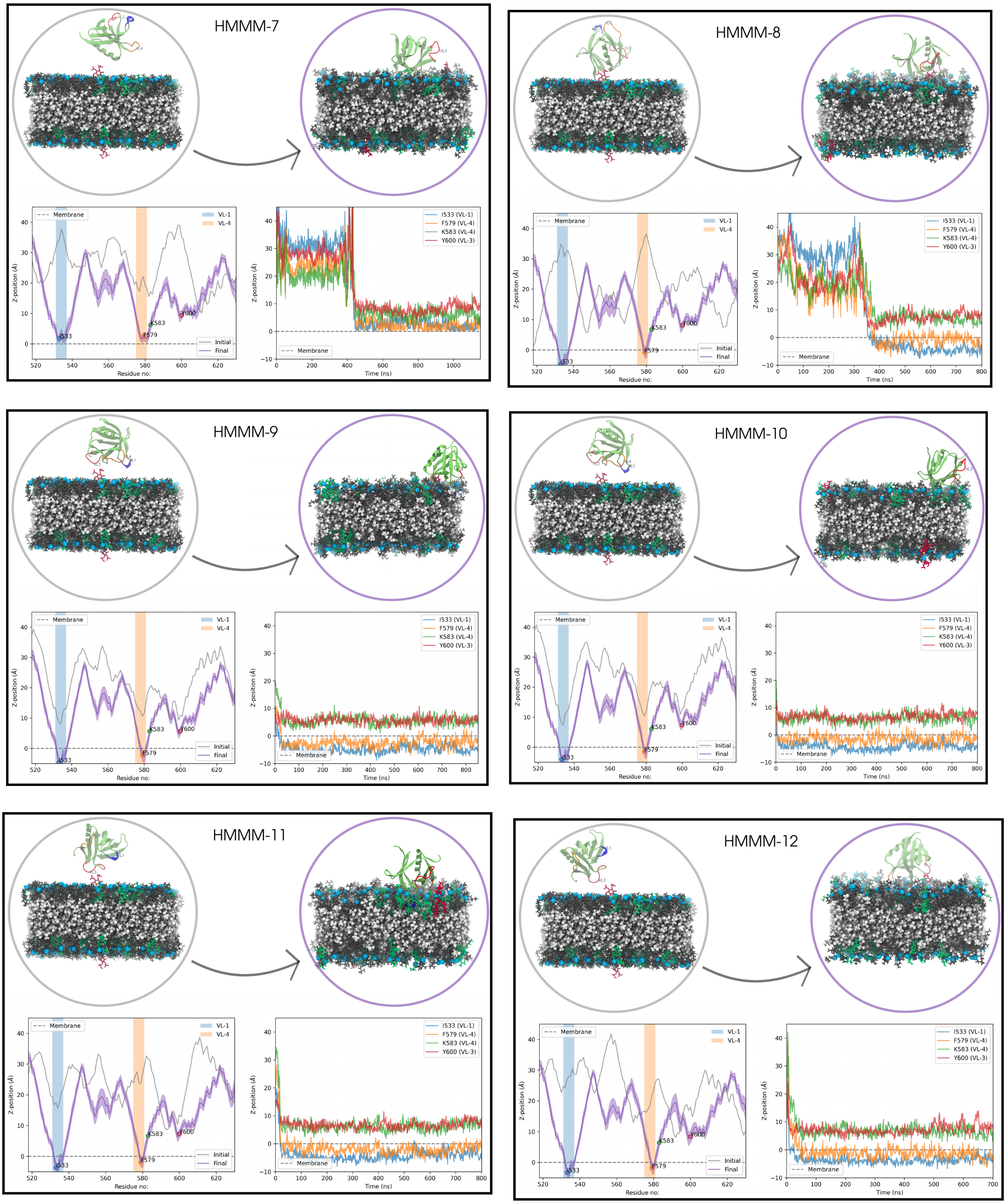


**Figure S4**: HMMM data for 12 replicates.

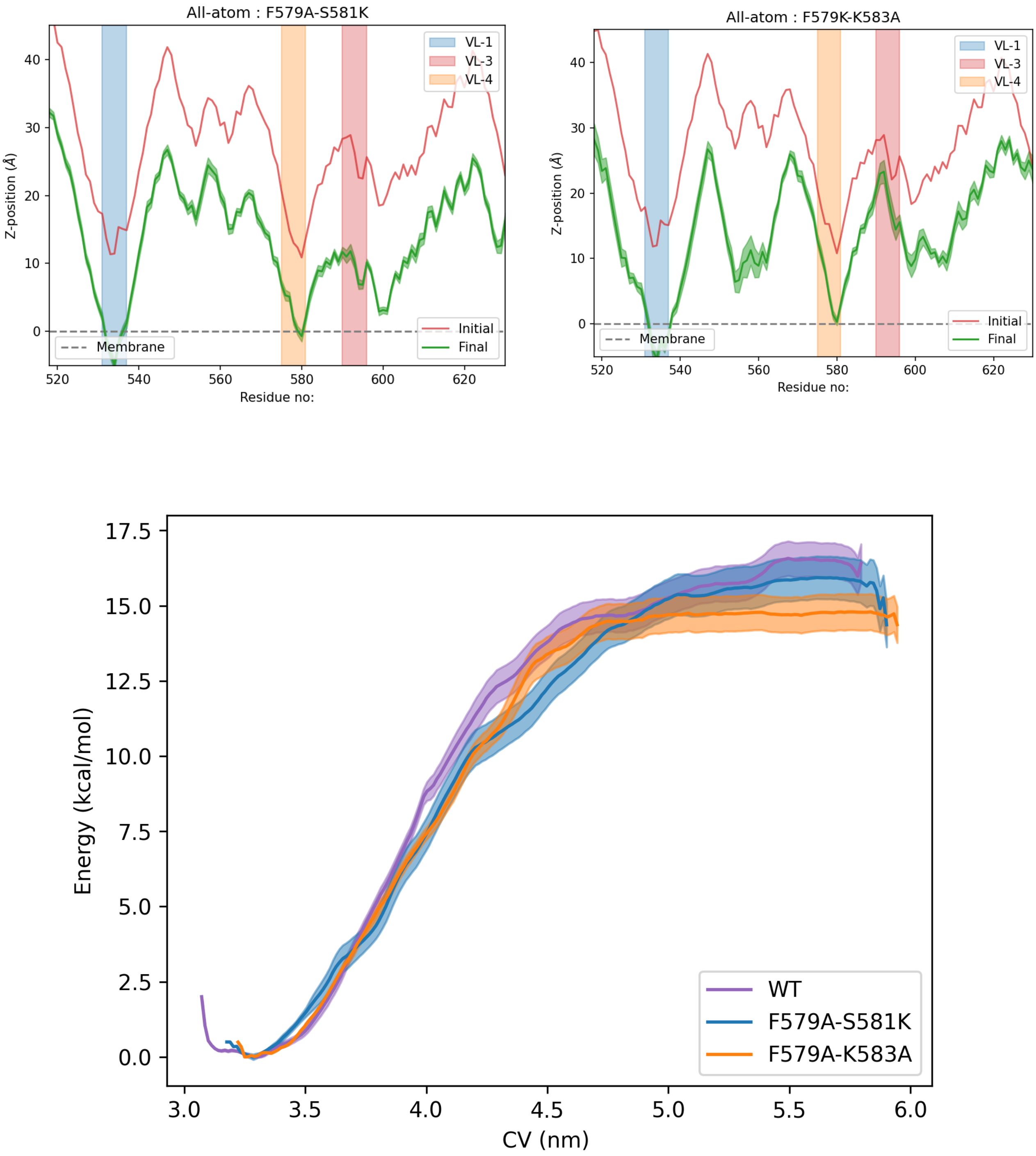


**Figure S5**: z-distance plots and US PMF data for double mutants

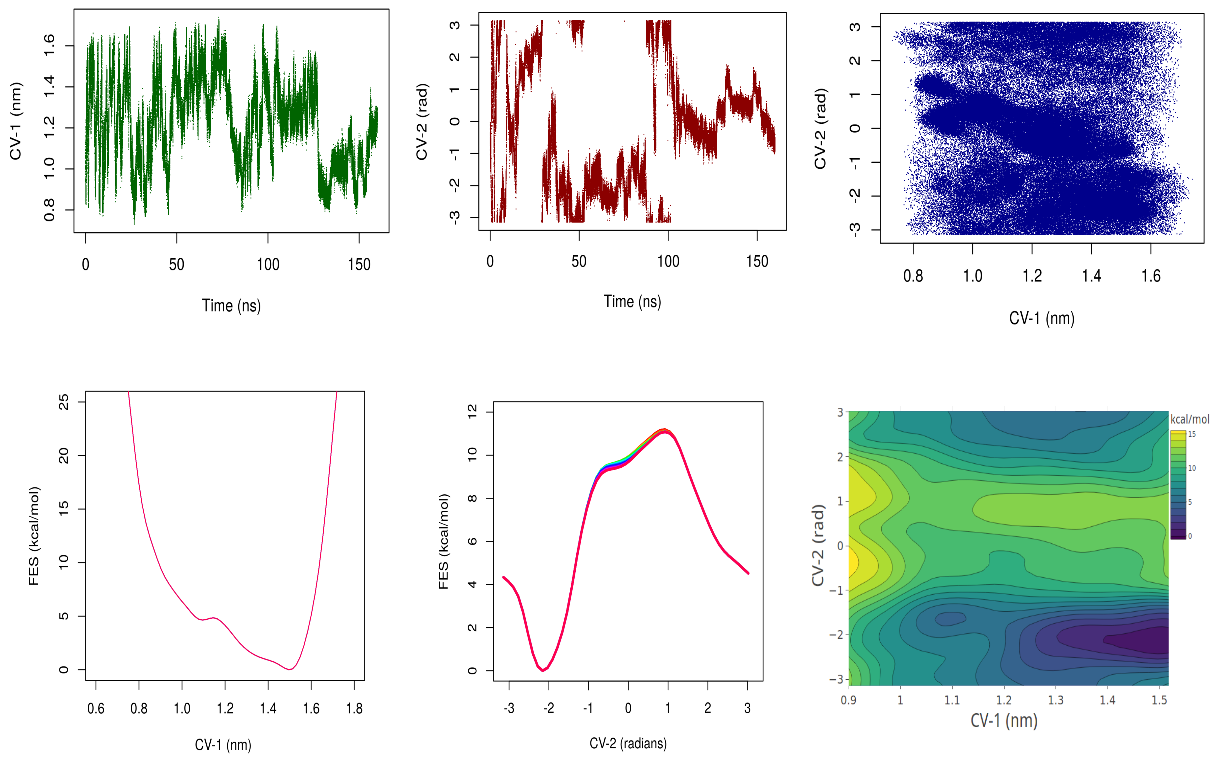
CV-1 (nm)
Time (ns)
CV-2 (rad)
Time (ns)
CV-2 (rad)
CV-1 (nm)
FES (kcal/mol)
CV-1 (nm)
FES (kcal/mol)
CV-2 (radians)
kcal/mol
CV-2 (rad)
CV-1 (nm)

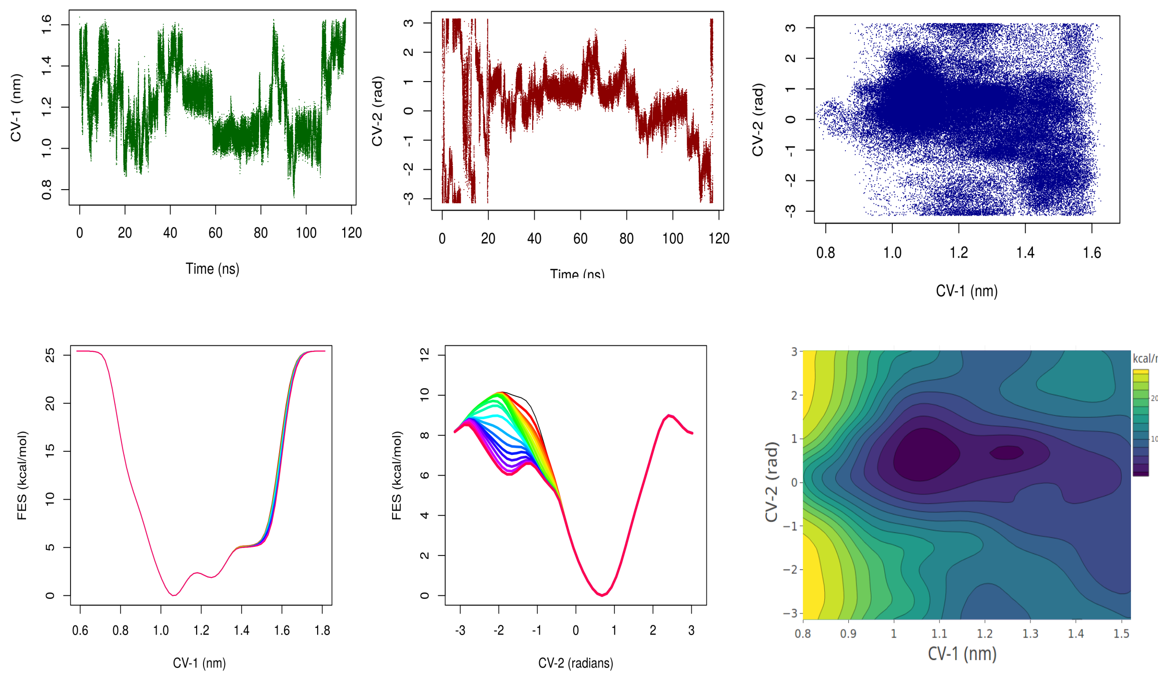
CV-1 (nm)
Time (ns)
CV-2 (rad)
Time (ns)
CV-2 (rad)
CV-1 (nm)
FES (kcal/mol)
CV-1 (nm)
FES (kcal/mol)
CV-2 (radians)
kcal/mol
CV-2 (rad)
CV-1 (nm)

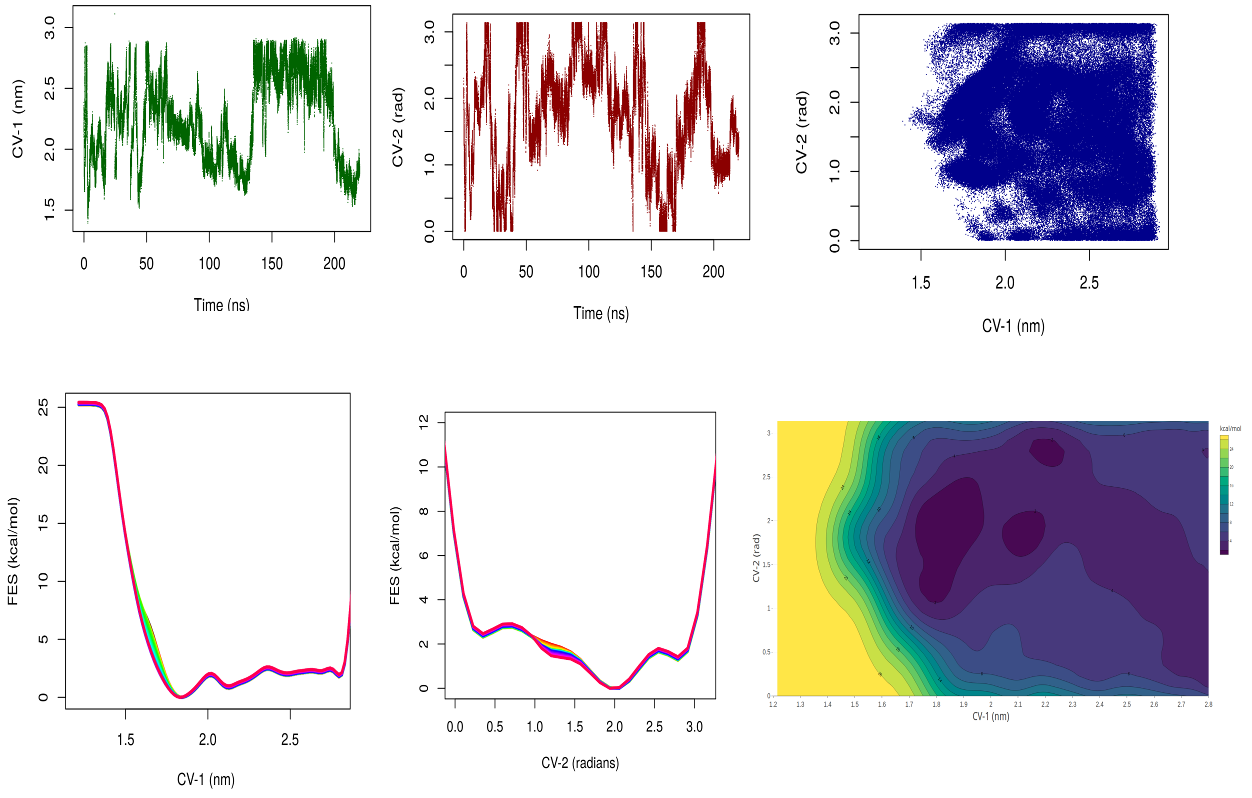


**Figure S6**: Convergence analysis for the well-tempered metadynamics runs. Top panel shows convergence results for simulations where with VL1-VL3 was chosen as the preferred docking area, middle panel is for system with VL1-VL4 as the preferred pocket and the bottom most panel is for system with no bias for any pocket. Top row for each panel shows the time evolution (sampling) of individual CV as well as both together. The bottom row for each panel shows the free energy surface (FES) data for individual CVs as well as when put together.

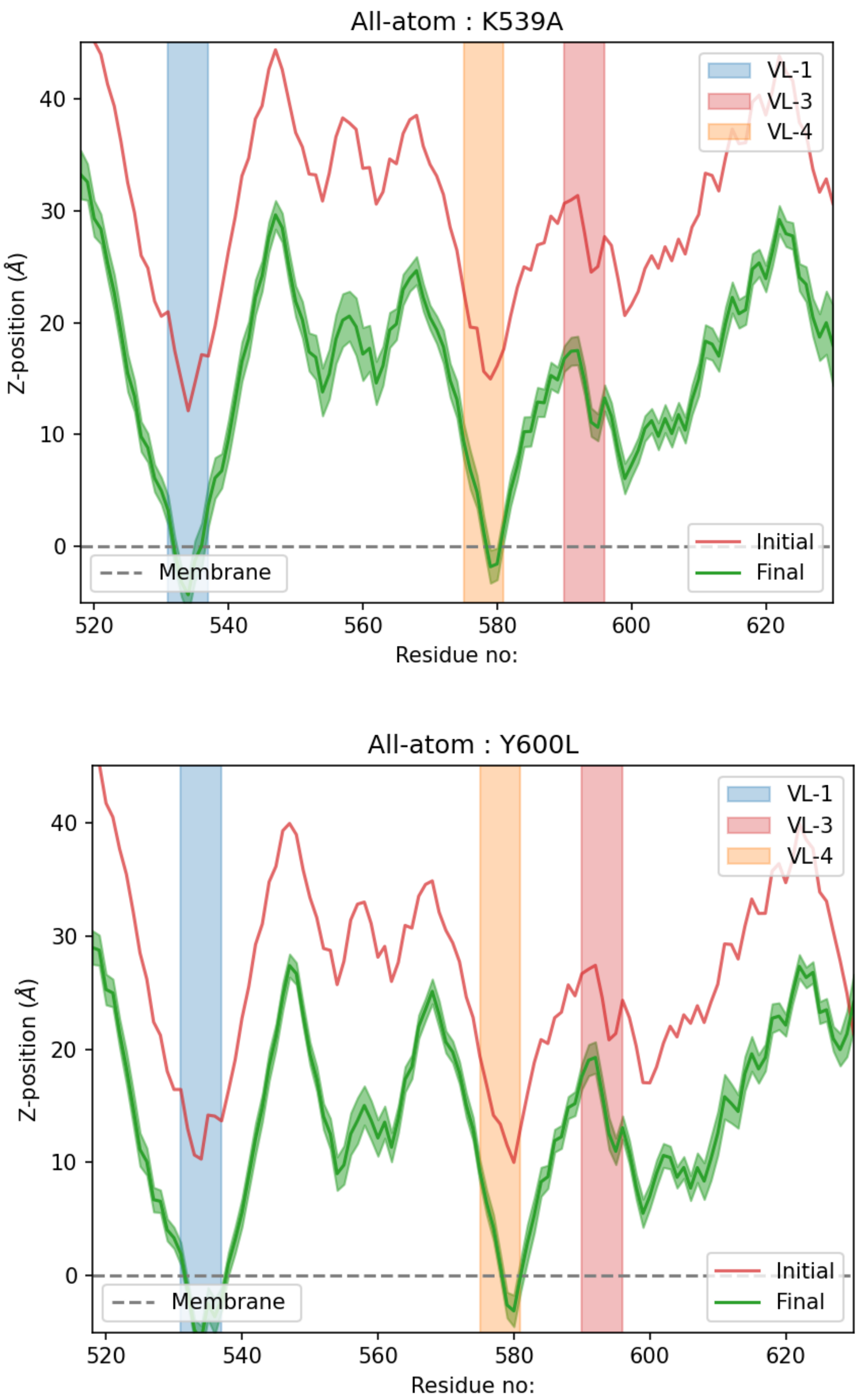


**Figure S7:** Z-distance analysis for K583A and Y600L from AAMD simulations

### 1. Well-Tempered Metadynamics Simulation Parameters:

-------------------------------------------------------------------

**1a. System: Loops 1-3 Well-Tempered metadynamics**

-------------------------------------------------------------------

Protein: 1DYN (Pleckstrin homology domain)
Ligand: Inositol triphosphate

**Definition of CVs:**

CV1: Distance between centre of mass of ligand's inositol ring and C-alpha atom of residue TYR 600 CV2: Torsion angle (A-B-C-D)

A: C-alpha atom of tyrosine-600
B: C-beta atom of tyrosine-600
C: C2 of inositol ring of ITP
D: C5 of inositol ring of ITP

**Metadynamics Parameters:**

Initial Height of gaussian: 1.0 kJ/mol

Guassian deposition rate: 1 ps

Length of the simulation: 160 ns
sigma of gaussian in CV1 direction:0.05 nm
sigma of gaussian in CV2 direction:0.35 rad

CV1 upper restraint: 1.5 nm with harmonic restraint whose kappa=3000

-------------------------------------------------------------------

**2b. System: Loops 1-4 Well-Tempered metadynamics**

-------------------------------------------------------------------

Protein: 1DYN (Pleckstrin homology domain)
Ligand: Inositol triphosphate

**Definition of CVs:**
CV1: Distance between centre of mass of ligand's inositol ring and C-alpha atom of residue 586 (phenylalanine)
CV2: Torsion angle (A-B-C-D)

A: N atom of phenylalanine 586
B: C-alpha atom of phenylalanine 586
C: C2 of inositol ring of ITP
D: C5 of inositol ring of ITP

**Metadynamics Parameters:**
Initial Height of gaussian: 1.0 kJ/mol

Guassian deposition rate: 1 ps

Length of the simulation: 110 ns
sigma of gaussian in CV1 direction:0.05 nm
sigma of gaussian in CV2 direction:0.35 rad

CV1 upper restraint: 1.5 nm with harmonic restraint whose kappa=3000

**-------------------------------------------------------------------**
**2c. System: Whole Protein Well-Tempered metadynamics**
**-------------------------------------------------------------------**

Protein: 1DYN (Pleckstrin homology domain)
Ligand: Inositol triphosphate

**Definition of CVs:**
CV1: distance between the centroid of protein and the centroid of ligand's inositol ring

CV2: Angle (A-B-C)
A: Centroid of the protein
B: C1 of inositol ring of ITP
C: C4 of inositol ring of ITP

**Metadynamics Parameters:**
Initial Height of gaussian: 1.0 kJ/mol
Guassian deposition rate: 1 ps

Length of the simulation: 220 ns
sigma of gaussian in CV1 direction:0.05 nm
sigma of gaussian in CV2 direction:0.35 rad

CV1 lower restraint: 1.2 nm with harmonic restraint whose kappa=2000
CV1 upper restraint: 2.8 nm with harmonic restraint whose kappa=2000

**2. Justification for using PHD instead of dynamin:** It can be argued that the deductions made from simulating the dyn1-PHD on the membrane might be limited in terms of transferability to what could be observed with the full dynamin structure. Though there would be changes, the choice can be justified by looking at some of the experimental work with PHD-mutant or knockouts. In Dar *et al* (1), PHD was replaced by a six residues long His-tag, which affects the fission rate but did not alter any other mechanism of dynamin. Dynamin formed a helix in a similar fashion as it does with WT-PHD. Though tightly coupled to each other, each domain of dynamin has a specific job assigned to it as is the case with most of the multi-domain proteins. Moreover, the sheer size of dynamin makes the computational work prohibitive, even with the largest of computational resource.

**3. Effect on bending modulus of membrane due to peripheral protein:** Recent article by Fowler *et al* (2) suggests that the contribution of peripheral protein in generating thermal undulation may not be significant enough to affect the bending modulus. The deductions were mostly based on simulations using a small protein tN-Ras. Our results turn out in contrast to the arguments made by the article. Our results show that the shallow-docking peripheral PHDs lower the bending modulus and here could be a couple of possible reasons that could explain this contradiction including the absence of cholesterol which is known to act as a knob to maintain packing "homeostasis" in the membrane. Another crucial factor that supports PHD association is the presence of $PIP_2$ in our lipid compositions. We observed in our CG simulations (with no $PIP_2$ molecule in their lipidome), that the PHDs don't stay close to the bilayer for the entire time and some of them actually diffuse long way from headgroup surface. Such behavior was not observed when $PIP_2$ was added to the system due to the specific binding that occurs between PHD and $PIP_2$. The other reason could be could be the nature of the protein itself. We would also like to discuss the reduction in bending modulus data in light of several recent studies where different kind of changes in membrane rigidity is observed due to peptides, peripheral and intrinsic proteins. There are several studies where peptides are shown to soften the membrane (3-6) and some very interesting recent work where certain peptides (GWALP in this case) could lead to a liquid-disordered system to have a higher rigidity than ordered systems (7).
